# Nuclear lamin B is crucial to the nuclear envelope integrity and extracellular trap release in neutrophils

**DOI:** 10.1101/647529

**Authors:** Yubin Li, Victoria P. Werth, Moritz Mall, Ming-Lin Liu

## Abstract

It’s not clear how nuclear envelope (NE) is ruptured for chromatin externalization during NETosis. The membrane rupture during neutrophil NET release was described as a membrane lysis process, this notion, however, has been questioned. Here, we found that lamin B, the structural NE component, was involved in NETosis. Unexpectedly, lamin B was not fragmented by destructive proteolysis, but rather disassembled into its intact full-length molecule, in NETotic cells with ruptured NE. In the mechanistic study, our experiments demonstrated that cytosolic PKCα translocated to the nucleus, where it serves as a NETotic lamin kinase to induce lamin B phosphorylation, following by lamina disassembly and NE rupture. To determine causality, we found that decreasing lamin B phosphorylation, by PKCα inhibition or genetic deletion, or mutation at the PKCα consensus phosphorylation sites of lamin B, attenuated extracellular trap formation. Importantly, strengthening NE by lamin B overexpression attenuated neutrophil NETosis *in vivo* and alleviated exhibition of NET-associated inflammatory cytokines in UVB irradiated skin of lamin B transgenic mice. These findings advance our understanding of NETosis process and elucidate a cellular mechanism that PKCα-mediated lamin B phosphorylation drives nuclear envelope rupture for NET release in neutrophils.

**Graphical Abstract:** 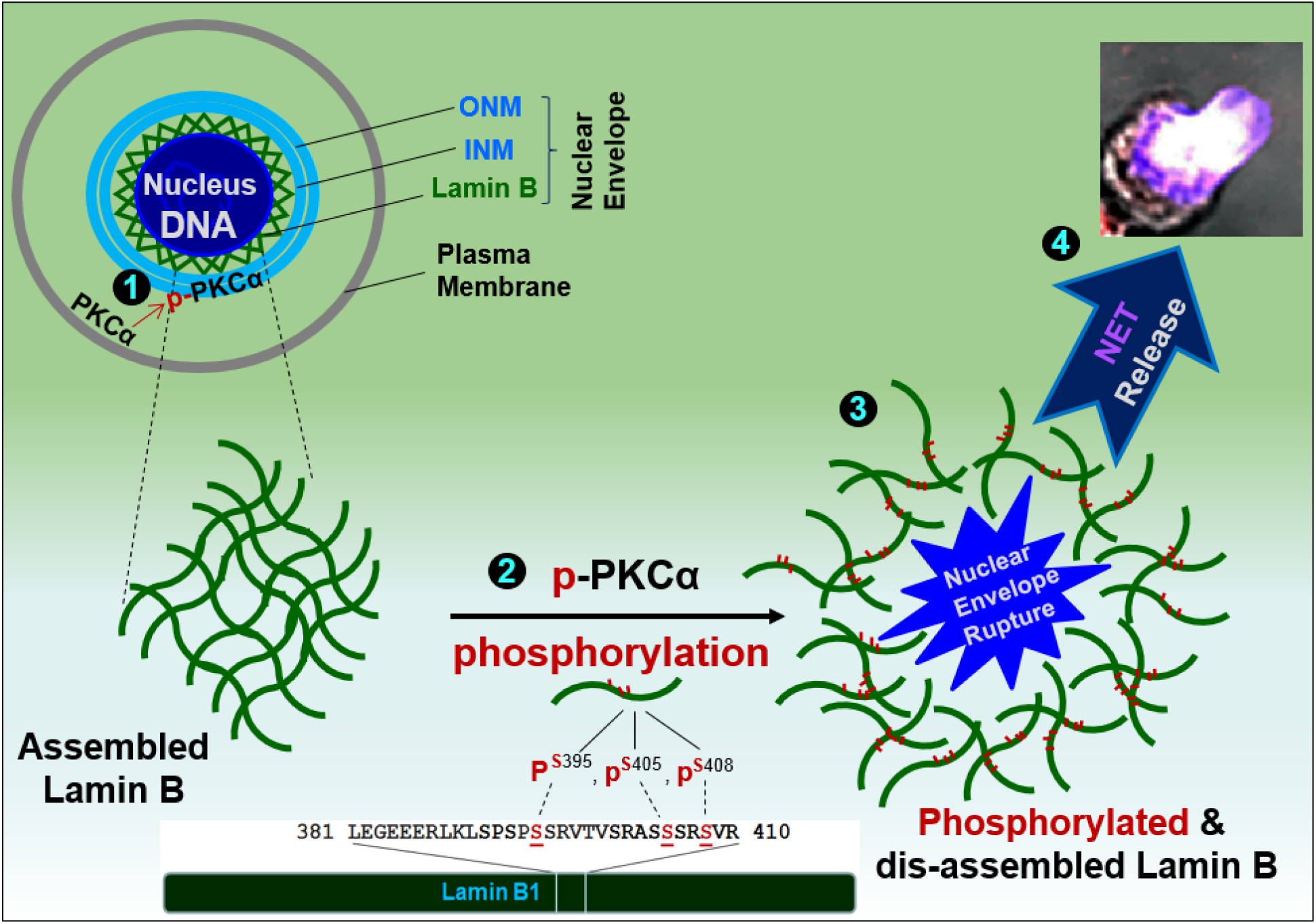

## INTRODUCTION

Neutrophils are the most common leukocytes. Increasing evidence from many studies (Brinkmann et al., 2004; Lood et al., 2016; Mayadas et al., 2014), including our own (Folkesson et al., 2015), indicates the importance of neutrophils in acute or chronic inflammation in variety of human diseases (Brinkmann et al., 2004; Jorch and Kubes, 2017; Mayadas et al., 2014; Papayannopoulos, 2017; Soehnlein et al., 2017). NETosis is a unique type of programmed neutrophil cell death with release of neutrophil extracellular traps (NETs) (Brinkmann et al., 2004). NETosis is involved in the pathogenesis of infectious (Brinkmann et al., 2004), athero-thrombotic (Warnatsch et al., 2015), metabolic (Wong et al., 2015), and autoimmune (Lood et al., 2016) diseases, as well as cancer (Jorch and Kubes, 2017).

NETosis is characterized by nuclear chromatin decondensation through peptidyl arginine deiminase type IV (PAD4) catalyzed histone citrullination (Neeli et al., 2008; Wang et al., 2009; Wong and Wagner, 2018), followed by chromatin extrusion via ruptured nuclear envelope. The externalized nuclear chromatin further mixes with cytosolic contents and granule proteins in the cytoplasmic compartment, and finally releases as NETs in extracellular milieu (Fuchs et al., 2007; Jorch and Kubes, 2017). Several signaling pathways may be involved in NETosis, i.e., production of reactive oxygen species (ROS) through NADPH oxidase (NOX2) dependent (Fuchs et al., 2007) or independent pathways (Douda et al., 2015), activation of protein kinase C (PKC), extracellular-signal-regulated kinase (ERK/MAPK), and phosphatidylinositide 3-kinase (PI3K)/Akt (Douda et al., 2014; Hakkim et al., 2011). Neutrophil elastase may also be involved in the process, including chromatin decondensation (Fuchs et al., 2007; Papayannopoulos et al., 2010), probably by neutrophil elastase-mediated histone degradation (Dhaenens et al., 2014; Papayannopoulos et al., 2010), while neutrophil elastase is dispensable for NET formation as neutrophils from neutrophil elastase deficient mice could still undergo NETosis in a recent study (Martinod et al., 2016).

Release of the decondensed nuclear chromatin, through the broken nuclear and plasma membranes, is the key step for formation of the neutrophil NET backbone. Neutrophil NET release was described as a lytic cell death mechanism (Fuchs et al., 2007; Papayannopoulos et al., 2010; Yipp and Kubes, 2013). However, this notion has been questioned by several recent observations (Pilsczek et al., 2010;Amulic et al., 2017; Neubert et al., 2018). Rupture of the nuclear envelope/nuclear membrane appears to be a distinct process from the previously described lysis or dissolution of the nuclear envelope (Amulic et al., 2017; Neubert et al., 2018). Since nuclear chromatin is enclosed by the nuclear envelope, its breakdown is, therefore, a prerequisite, and a key step, for externalization of nuclear chromatin and extracellular trap formation. (Papayannopoulos et al., 2010; Yipp and Kubes, 2013) However, the relevant mechanisms that regulate nuclear envelope rupture in NETosis are still unclear.

The nuclear envelope consists of outer and inner lipid nuclear membranes (ONM and INM) and nuclear lamina, a protein filament meshwork that provides structural scaffold to reinforce nuclear envelope integrity (Goldberg et al., 2008; Schreiber and Kennedy, 2013; Stewart et al., 2007). Nuclear lamins are categorized as either A-type (A, C) or B-type (B_1_, B_2_) lamins (Schreiber and Kennedy, 2013). A-type lamins form thick filament bundles that provide mechanical rigidity to the nucleus (Goldberg et al., 2008; Lammerding et al., 2006; Rowat et al., 2013), whereas B-type lamins form thin but highly organized meshworks that are crucial to the integrity of the nuclear envelope (Goldberg et al., 2008; Vergnes et al., 2004). Lamin B is anchored underneath the INM through lamin B receptor (LBR) (Schreiber and Kennedy, 2013; Stewart et al., 2007). Nuclear envelope breakdown (Hatch and Hetzer, 2014) is a common cellular event in nuclear fragmentation during cell apoptosis (Shimizu et al., 1998), in nuclear division during cell mitosis (Collas et al., 1997; Mall et al., 2012), and in viral nuclear access during viral infection (Park and Baines, 2006). In the above cellular processes, the nuclear lamin B is either proteolytically degraded by caspases (Cross et al., 2000), or disassembled through phosphorylation by protein kinase C (PKC) (Collas et al., 1997; Fay and Pante, 2015; Muranyi et al., 2002).

Mature neutrophils are terminally differentiated cells and were thought to have an unusual nucleus with a paucity of lamin A/C (Olins et al., 2008) and a reduced amount of lamin B (Olins et al., 2008; Wong et al., 2013). Recent studies, however, reported that neutrophils do have A-type (Amulic et al., 2017) and B-type (Moisan and Girard, 2006; Rowat et al., 2013) lamins. The role of lamin B in neutrophil biology is unclear, and the involvement of lamin B in neutrophil NETosis has not been investigated. In the current study, we investigated the importance of lamin B in nuclear envelope integrity in neutrophils and cellular mechanisms that regulate nuclear envelope rupture during NETosis in neutrophils from human and mice. We also investigated the effect of strengthening nuclear envelope by lamin B overexpression on neutrophil NETosis *in vivo* and exhibition of NET-associated inflammatory cytokines in the UVB-irradiated skin of lamin B transgenic mice.

## RESULTS

### Nuclear lamin B is a substantial component of nuclear envelope that is involved in neutrophil NETosis

Neutrophils are terminally differentiated cells, and the role of nuclear lamina in neutrophil biology is not well understood (Olins et al., 2008). However, nuclear envelope breakdown is required for nuclear chromatin externalization during NET formation. To explore the involvement of lamin B, we detected lamin B expression in human primary pPMNs (Fig. 1A,B, Fig. S1A), mouse primary mPMNs (Fig. S1B), and human neutrophil-like dPMNs (Fig. S1C), by immunoblotting or confocal microscopy. In the HL-60 derived dPMNs, lamin B expression was comparable throughout the 5-day neutrophil differentiation period (Fig. S1C). Neutrophil NETosis can be triggered by PMA, the most commonly used NET inducer (Fig. 1A) (Brinkmann et al., 2004), or by platelet activating factor (PAF), a UVB-induced proinflammatory lipid mediator and naturally existing stimulus of neutrophil NETs (Fig. 1E-G) (Damiani and Ullrich, 2016). Confocal microscopy analyses indicated that nuclear lamin B is a substantial component of the nuclear envelope (Fig. 1B-C), which is ruptured during neutrophil NETosis (Fig. 1B), indicating the involvement of lamin B in the process of neutrophil NETosis.

**Figure 1.**
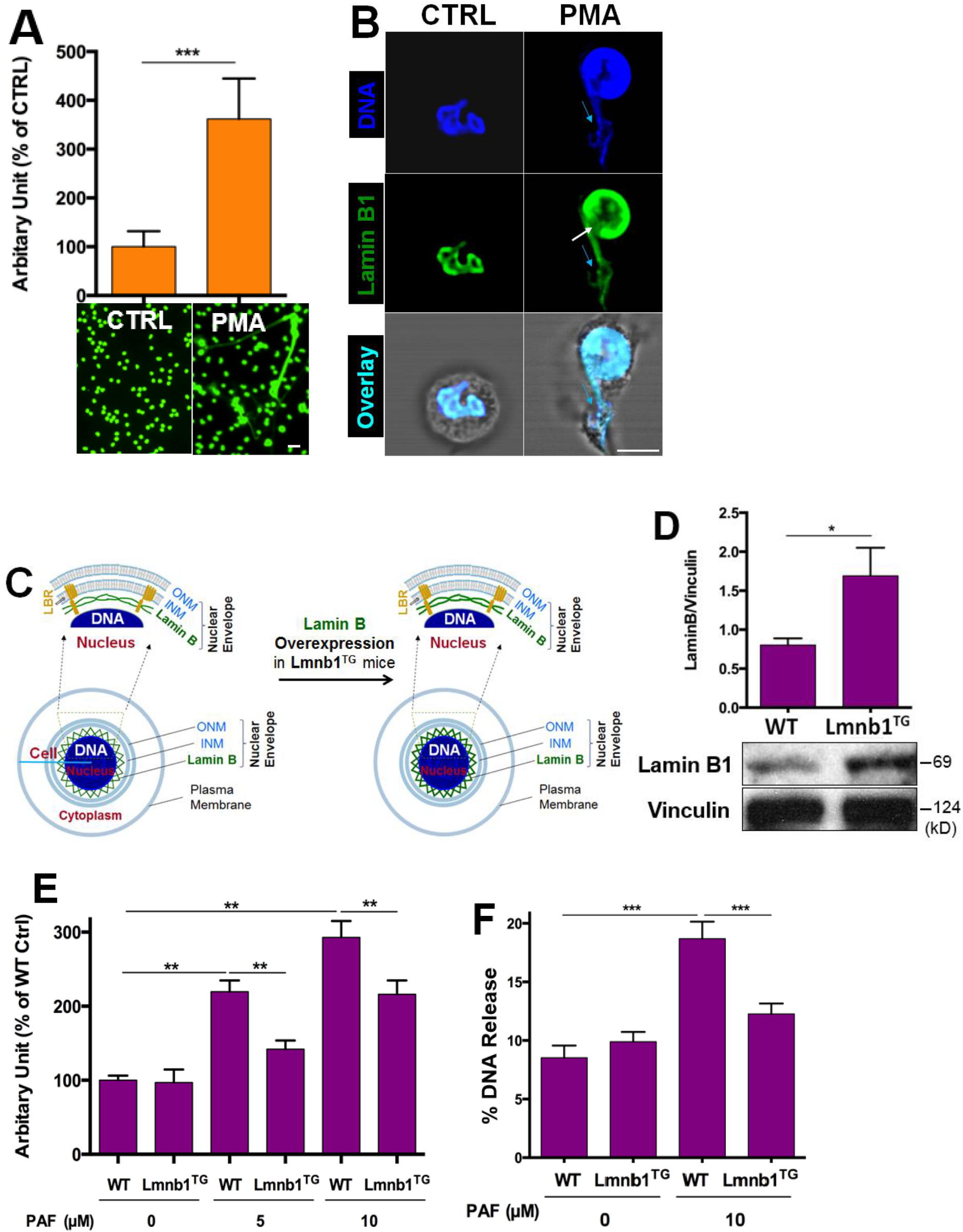
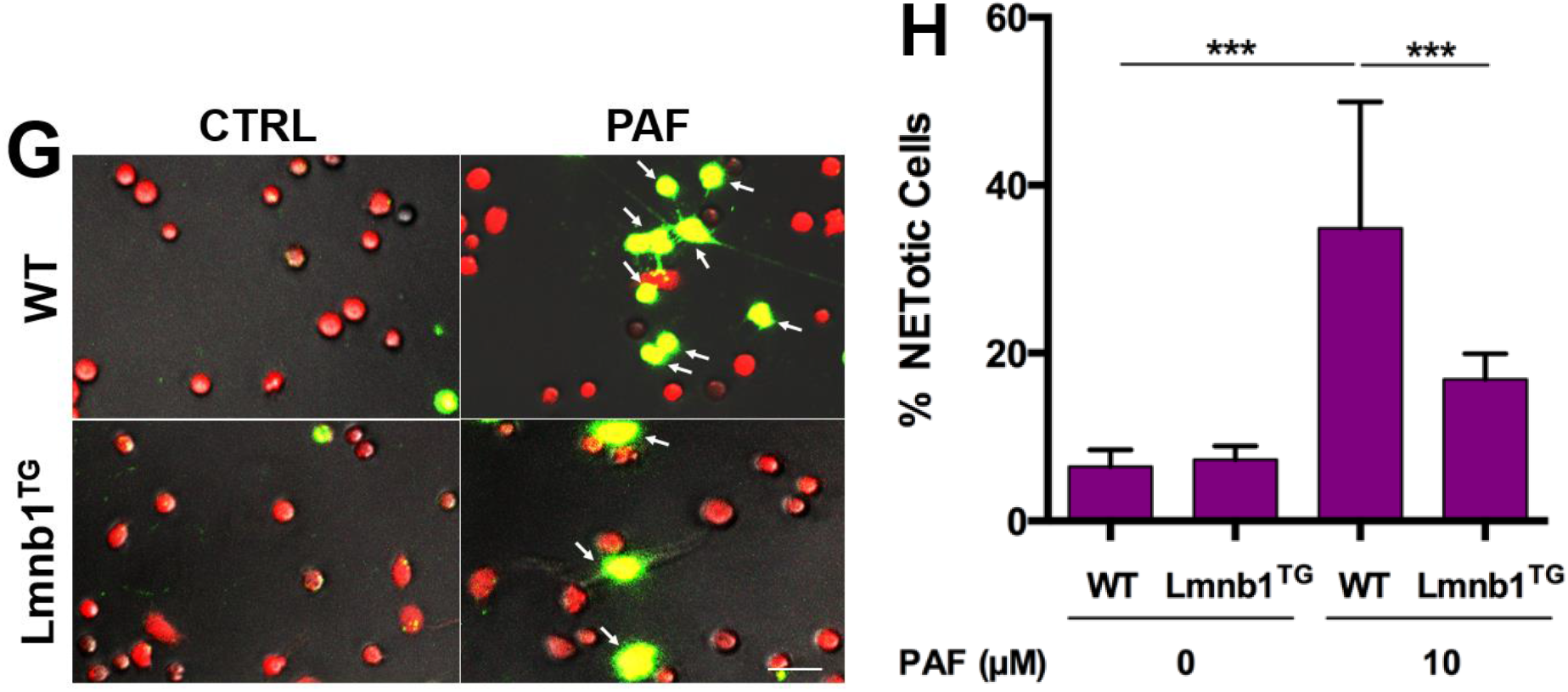
Nuclear lamin B is a substantial component of nuclear envelope that is involved in NET release in neutrophils. (**A**) The summary and representative analysis of NETosis of pPMNs stimulated by 50 nM PMA for 3h, detected by fluorescent microplate reader and confocal microscopy, respectively. (**B**) Confocal microscopy analysis of the representative pPMNs that were treated without (control) or with 50 nM PMA for 3h, and then stained for lamin B and DNA as described in panel B. The white arrow demonstrates the site of nuclear envelope rupture, the light blue arrows indicate the release of decondensed DNA associated with lamin B molecules. Scale bars, 40 µm (A) 10 µm (B) respectively. (**C**) Schematic cross-section of a cell and portion of the nucleus and the nuclear envelope, as well as the ones with corresponding overexpression of lamin B. The two lipid bilayers of the nuclear envelope are the inner and outer nuclear membranes (INM and ONM, respectively). The meshwork, nuclear lamin B is anchored to INM through lamin B receptor (LBR). (**D**) Representative and summary of immunoblots of lamin B1 of bone marrow mPMN neutrophils from WT and Lmnb1^TG^ mice. Vinculin served as loading control. (**E**) Summary analysis of PAF-induced NETosis in mPMNs from WT vs Lmnb1^TG^ mice that were stimulated without or with 10 µM PAF for 3h, then fixed by 2% PFA and stained by Sytox Green, following by detection with fluorescent microplate reader. (**F**) The endpoint analysis of NET-DNA release index was detected by coincubation of primary mouse peritoneal mPMNs from WT and Lmnb1^TG^ mice without (control) or with 10 µM PAF in medium containing 1 μM Sytox Green dye with recording by a microplate reader at the 3h time point. The NET-DNA release index was reported in comparison to an assigned value of 100% for the total DNA released by neutrophils lysed by 0.5% (v/v) Triton-X-100. (**G,H**) Representative images (G) and summary analysis (H) of PAF-induced NETosis in mPMNs from WT vs Lmnb1^TG^ mice that were stimulated without or with 10 µM PAF for 3h, and stained with both cell permeable Syto Red and cell impermeable Sytox Green, without fixation. Images were taken by Olympus confocal microscopy, followed by automated quantification of NETs on 5-6 non-overlapping area per well using ImageJ for calculation of % NETotic cells. (G) Scale bars, 40 μm. Panels A,D,E,F,H were summary analysis of NETosis (A,E), or summary analyses of immunoblots (D), DNA% release index (F) that were calculated based on the arbitrary fluorescent readout unit (A,E), immunoblot arbitrary unit (D), NET-DNA release index calculated based on fluorescent readout (F), or % NETotic cells by image analysis (H) from more than 3 independent experiments. P*<0.05, P**<0.01, P***<0.001 between groups as indicated. Comparisons amongst three or more groups were performed using ANOVA, followed by Student-Newman-Keuls test. Comparison between two groups was analyzed by student t test.

Lamin B is known to be assembled as a filament meshwork that lies beneath the INM, and the two are associated by an integral protein, lamin B receptor (Fig. 1C). Given the importance of lamin B in nuclear envelope organization (Goldberg et al., 2008; Vergnes et al., 2004), one may propose that lamin B may be important in regulation of nuclear envelope rupture and NET release in neutrophils. To test this hypothesis, we sought to explore the effects of lamin B overexpression on neutrophil NETosis by using neutrophils from lamin B1 transgenic Lmnb1^TG^ mice whose cells (Fig. 1D) have increased expression of lamin B. Interestingly, we found the attenuated NETosis in peritoneal neutrophils with elevated levels of nuclear lamin B1 expression from Lmnb1^TG^ mice, as compared to those from their WT littermates (Fig. 1E-G), analyzed by conventional fluorometric NET quantification (Chicca et al., 2018; Sollberger et al., 2016), the endpoint analysis of NET-DNA release index (Douda et al., 2015; Khan et al., 2018), and immunofluorescent imaging analysis of NETosis (Martinod et al., 2017; Sollberger et al., 2016). On the other hand, it is known that lamin B maturation is regulated by farnesylation (Adam et al., 2013), and long-term inhibition of protein farnesylation with farnesyltransferase inhibitor (FTI) can reduce the amount of mature lamin B which affects nuclear function in several cell types (Adam et al., 2013). To explore the effects of reduced lamin B, we found that co-incubation with FTI can reduce the amount of mature lamin B in RAW 264.7 (Fig. S1D). Most importantly, our results showed that the reduced amount of lamin B in FTI-pretreated cells could enhance PAF-induced extracellular trap formation (Fig. S1E-F). Taken together, the levels of nuclear lamin B expression affect extracellular trap formation in a negative dose-dependent manner.

Therefore, our results demonstrated that nuclear lamin B is a substantial component of the nuclear envelope in neutrophils, similar to other cell-types (Goldberg et al., 2008; Vergnes et al., 2004). Our study indicates that nuclear lamin B is crucial to nuclear envelope integrity during neutrophil NET release.

### Nuclear lamin B disassembly, but not proteolytic cleavage, is responsible for NETotic nuclear envelope rupture in neutrophils

Next, we sought to elucidate how lamin B is involved in nuclear envelope rupture during NETosis. Unexpectedly, the immunoblot analysis, of the time-course studies of human dPMNs and primary pPMNs using whole-cell lysates, demonstrated that the nuclear lamin B remained as an intact full-length molecule over the 3h of neutrophil NETosis induction upon stimulation by PMA (Fig. 2A-B) or PAF (Fig. S2A-B), in contrast to the destructively cleaved and fragmented lamin B in apoptotic neutrophils (Fig. 2A). Importantly, immunoblot analysis of the released extracellular NETs (Fig. 2C), isolated from conditioned medium of NETotic pPMNs, also showed the nuclear lamin B released with NETs to the extracellular space remaining as a full-length molecule (Fig. 2C), which confirmed the findings from the nuclear lamin B of the whole-cell lysates. Therefore, our results suggested the potential lamin B disassembly, but not the destructive cleavage, is responsible for the NETotic NE rupture in neutrophils.

**Figure 2.**
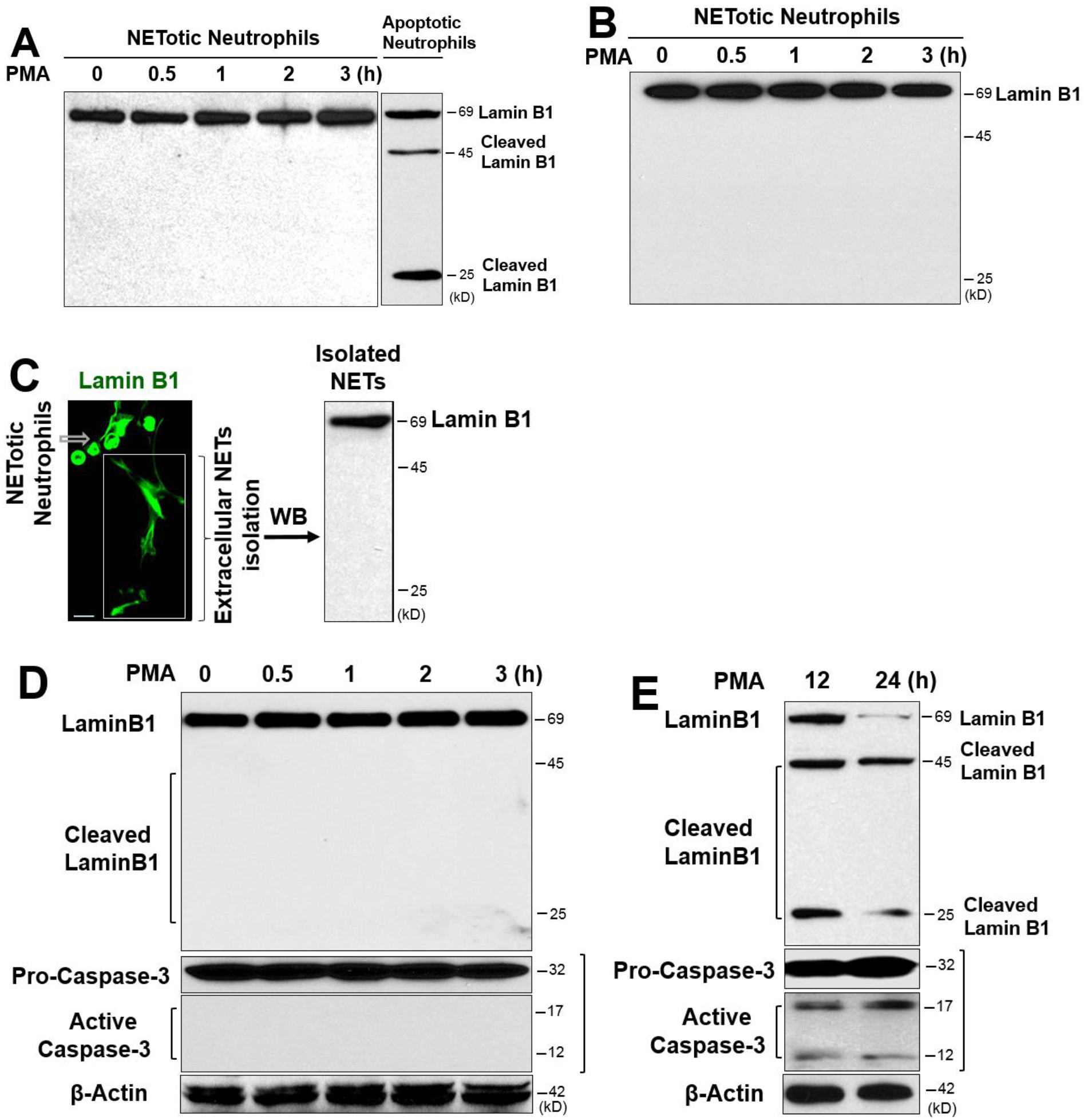
Nuclear lamin B disassembly, but not proteolytic cleavage, is responsible for NETotic nuclear envelope rupture in neutrophils. (**A)** Representative immunoblot image of full-length lanes of lamin B1 in human dPMNs that were treated with PMA for 0, 0.5, 1, 2, 3h during NETosis induction, or apoptotic dPMNs that were induced by PMA for longer term (12h) treatment. (**B**) Representative immunoblots of lamin B1 in primary human pPMNs that were treated with PMA (B) for 0, 0.5, 1, 2, 3h during neutrophil NETosis. (**C**) Representative confocal microscopy image of a group of NETotic pPMNs and the extracellular NETs released by these cells, as well as the immunoblot analysis of the lamin B1 with the lysate of the NETs isolated from the conditioned medium of NETotic pPMNs that were induced by 3h PMA treatment. The arrow indicates NETotic neutrophils, the square demonstrates the extracellular NETs under confocal microscopy. (**D**) Representative immunoblots with full-length lanes for lamin B1 analysis, and pro-caspase-3 and its activated form caspase-3 in NETotic human dPMNs that were treated with PMA for 0, 0.5, 1, 2, 3h. (**E**) Representative immunoblots display full-length lamin B1 (69 kDa) and their cleaved fragments (45 and 25 kDa), and pro-caspase-3 and its activated form caspase-3 in the apoptotic human dPMNs that were treated with PMA for 12, 24h. The anti-human lamin B1 was used in all immunoblots (A-E) and the confocal image (C), and the latter was further detected by FITC-labeled 2^nd^ Ab, scale bar, 20 μm. In panel D, E, anti-human caspase-3 was used, and β-actin served as loading control.

Caspase-3 mediates apoptosis and cleaves lamin B (Slee et al., 2001) during apoptotic nuclear fragmentation. In the current study, we found that caspase-3 remained inactive as pro-caspase-3 over 3h period of time during NETosis induction (Fig. 2D), while activated caspase-3 was detected in apoptotic dPMNs that were caused by extended PMA treatment (Arroyo et al., 2002) for 12-24h (Fig. 2E). Furthermore, we saw that lamin B was cleaved into 25 and 45 KDa fragments, corresponding to caspase-3 activation during apoptosis (Fig. 2E). These experiments further confirmed that the destructive proteolytic cleavage is not responsible for lamin B disintegration during nuclear envelope rupture in NETotic neutrophils.

### Nuclear translocation and phosphorylation of PKCα during neutrophil NETosis

To address how nuclear lamin B was disassembled in NETotic nuclear envelope rupture, we found that cytosolic PKCα was translocated to nucleus where PKCα may mediate nuclear lamin B phosphorylation, disassembly, and nuclear disintegration, similar to the context of mitosis (Mall et al., 2012), and viral infection (Park and Baines, 2006). The time-course experiments (Fig. 3A) demonstrated that PKCα was diffusely distributed in the cytoplasm at time 0, but gradually translocated into the nucleus over the time during NETosis induction. PKCα accumulated in the nucleus at 60-min time point, and resulted in nuclear envelope discontinuity/ rupture (Fig. 3B), followed by the consequent extrusion of the decondensed chromatin from the ruptured nuclear envelope and the eventual collapse of the nuclear envelope at 180 min (Fig. 3A), and the subsequent release of neutrophil NETs in the extracellular space. Correspondingly, PKCα phosphorylation was gradually increased over the 3h period of time in the immunoblot analysis of human primary pPMNs (Fig. 3C,D) and dPMNs (Fig. S3A-B) stimulated by PMA or PAF.

**Figure 3.**
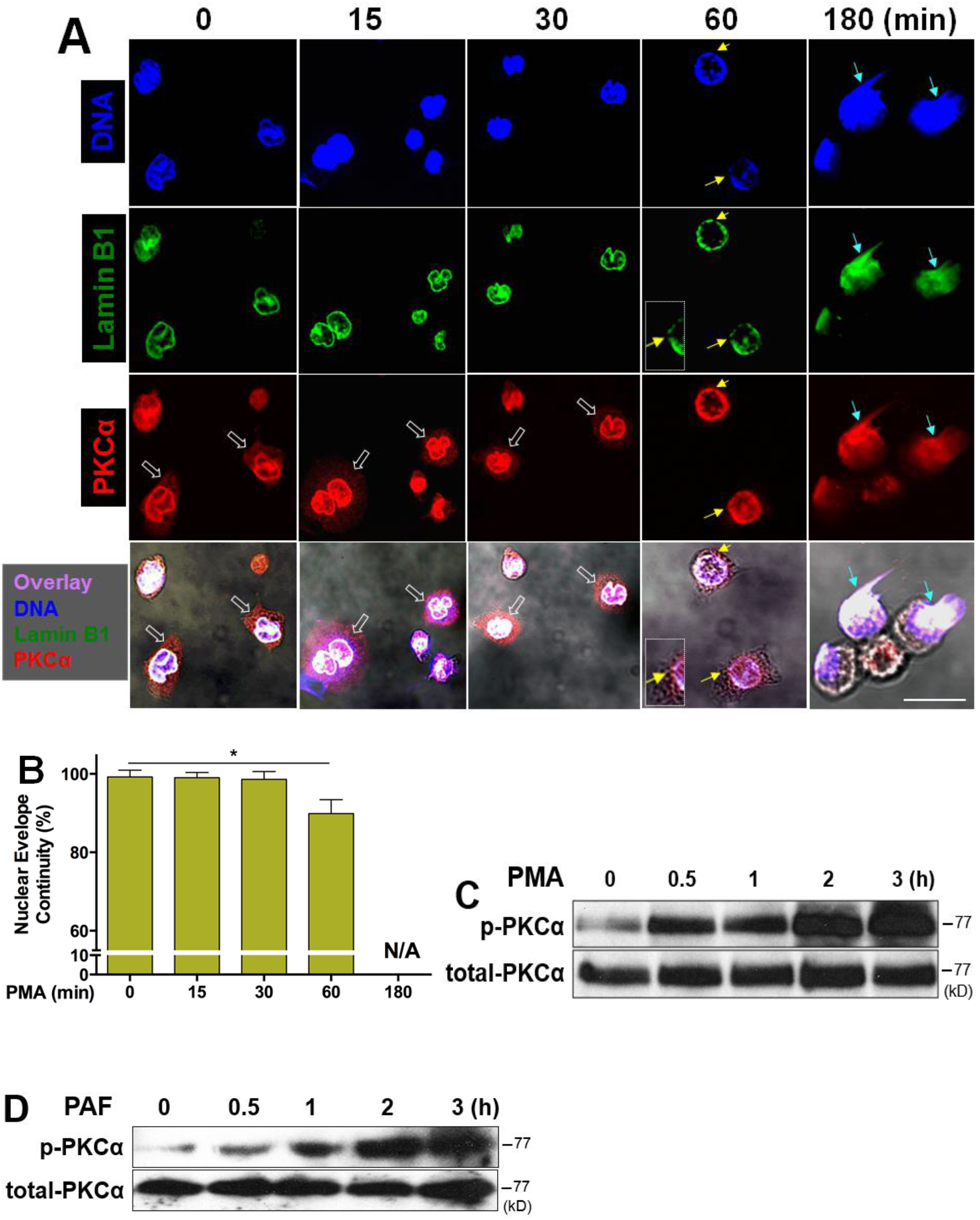
Nuclear translocation and phosphorylation of PKCα is accompanied by nuclear envelope rupture during neutrophil NETosis. (**A)** Representative images for the time course of PKCα nuclear translocation, subsequent nuclear envelope rupture, and DNA release in human dPMNs exposed to 50 nM PMA for 0, 15, 30, 60, and 180 min, then stained concomitantly for DNA **(DAPI)**, nuclear lamin B (primary anti-lamin B, and **FITC**-labeled secondary antibody), and PKCα (primary anti-human PKCα, and **PE**-labeled secondary antibody), following by confocal fluorescent microscopy analysis. White-empty arrows indicate cytoplasmic distribution of PKCα at 0, 15, and 30 min, yellow arrows indicate site of discontinuity/rupture of nuclear envelope at 60 min, light blue arrows display the sites of nuclear envelope rupture and chromatin release at 180 min (A). Scale bars, 20 μm. (**B)** Summary analysis of the nuclear envelope continuity was analyzed based on staining of the nuclear envelope with primary anti-lamin B, and **FITC**-labeled secondary antibody. (**C,D**) Representative immunoblots of total PKCα and p-PKCα in primary human pPMNs that were treated by PMA (C) or PAF (D) for 0, 0.5, 1, 2, 3h. The summary analyses of panel B were calculated based on the circumferences/perimeters of the cell nuclei of 7-10 cells from different time points from more than 3 independent experiments. P*<0.05 between groups as indicated. Comparisons amongst three or more groups were performed using ANOVA, followed by Student-Newman-Keuls test.

### Accumulation of phosphorylated PKCα in nucleus results in lamin B phosphorylation, disassembly, and nuclear envelope rupture during neutrophil NETosis

To explore the potential role of phosphorylated PKCα (p-PKCα) in nuclear envelope rupture in NETosis, we firstly isolated the nucleus of NETotic neutrophils, and found the accumulation of p-PKCα and the corresponding full-length lamin B in the nuclear fraction by immunoblots (Fig. 4A). The morphological analysis of NETotic neutrophils by confocal microscopy also confirmed the accumulation of p-PKCα at the site of nuclear envelope disruption (Fig. 4B) or rupture (Fig. 4C), and the release of the disassembled lamin B (Fig. 4B-D) with the decondensed chromatin (Fig. 4D) to form extracellular NETs in NETotic neutrophils. Furthermore, the p-PKCα and full-length lamin B1 were also detected in the extracellular NETs (Fig. 4E) isolated from cultured NETotic human primary pPMNs. The results of concomitant detection of these two molecules indicated a potential interaction between p-PKCα and lamin B in the nuclear envelope.

**Figure 4.**
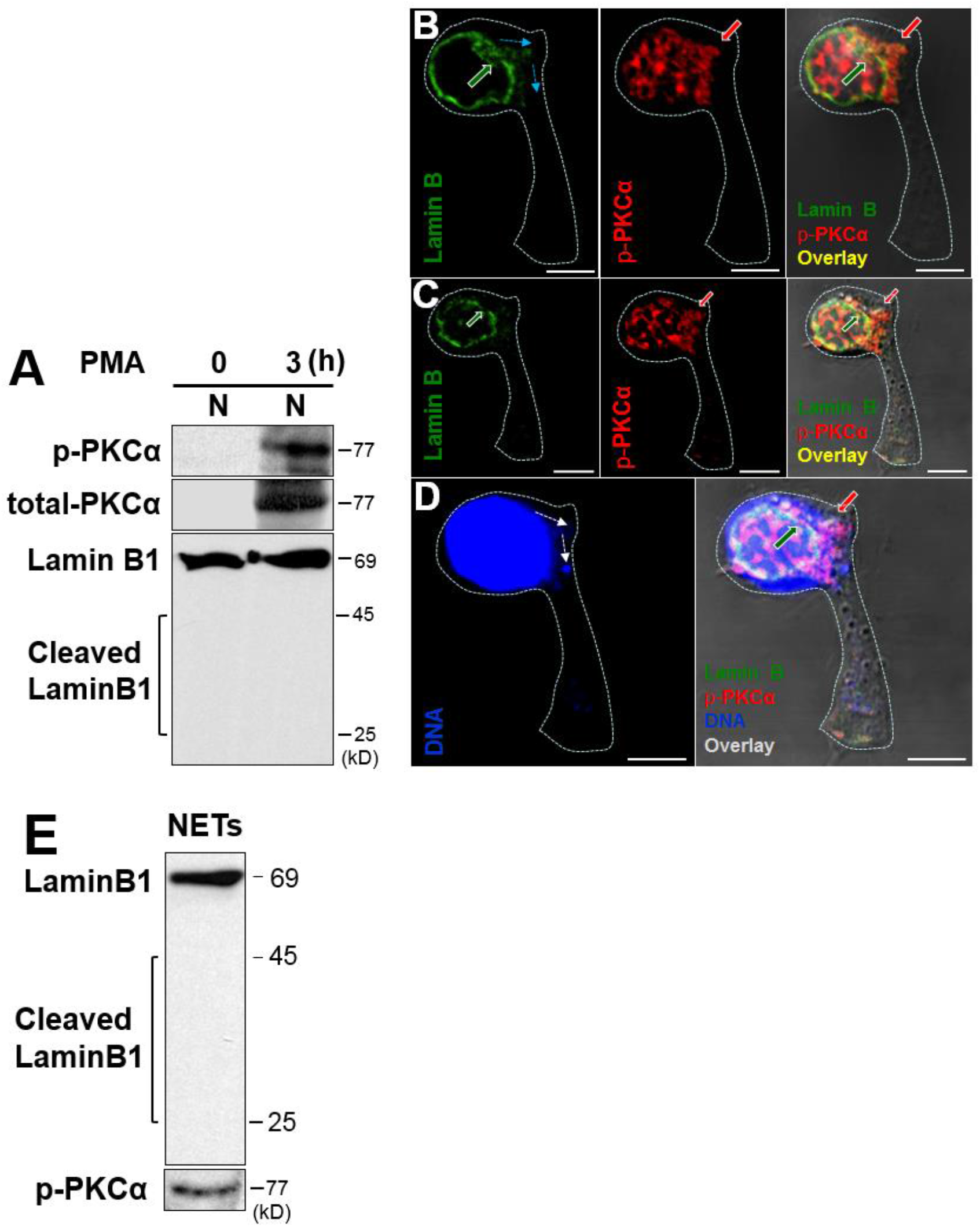
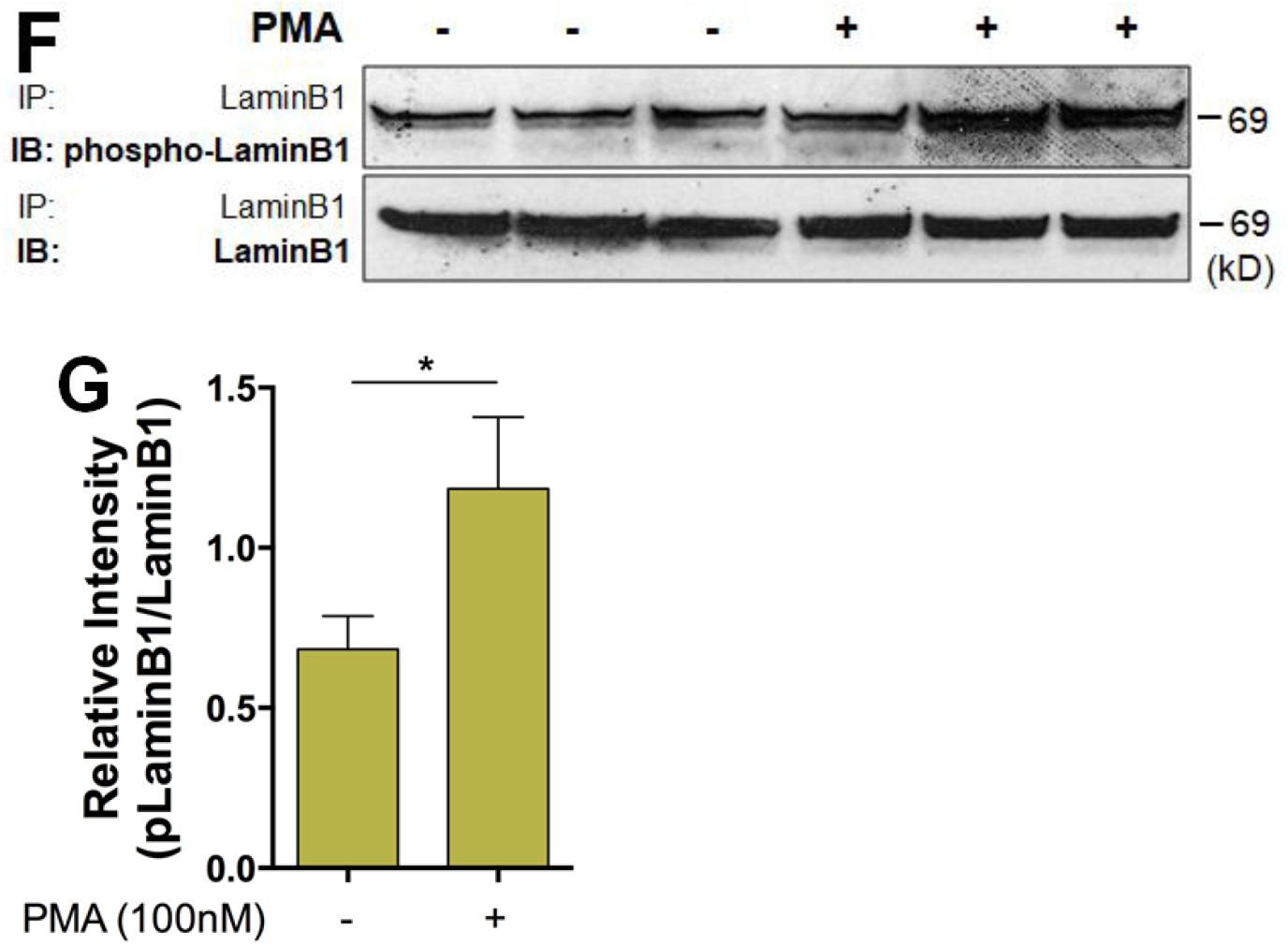
Accumulation of phosphorylated PKCα in nucleus results in nuclear lamin B phosphorylation, disassembly, and the consequent nuclear envelope rupture during neutrophil NETosis. (**A**) Representative immunoblots of p-PKCα and total PKCα of isolated nuclei and the corresponding status of lamin B1 in human dPMNs that were treated with PMA for 0 or 3h. (**B-D)** Representative sectional images of the nucleus of a NETotic neutrophil induced by PMA, stained for lamin B (primary anti-lamin B, and **FITC-**labeled secondary antibody), p-PKCα (primary anti-human p-PKCα^S657^, and **PE**-labeled secondary antibody), and nuclear DNA (**DAPI**). The sections on panels B-D indicate a NETotic neutrophil with a ruptured nuclear envelope (green arrows indicate the site of rupture), accumulation of p-PKCα (red arrows pointed), and release of the disassembled lamin B (light blue arrows with dash line) and de-condensed DNA (white arrows with dash line). Sections B to D were taken to visualize the different planes of the z stack sectioning progressively from the apical surface to their basal surface of the cells. Scale bars, 10µm. (**E**) Immunoblot images of lamin B1, then stripped and probed for p-PKCα, of the homogenates of NETs isolated from conditioned medium of NETotic primary human pPMNs that were treated by PMA. (**F,G**) Representative and summary immunoblot (IB) detection of phospho-lamin B and total lamin B with the lamin B protein purified by immunoprecipitation (IP) with anti-lamin B from human dPMNs that were treated either by PMA (**F,G**) for 0 or 3h. Comparison between two groups was analyzed by student t test.

In several other contexts of the nuclear envelope rupture (Kuga et al., 2010; Machowska et al., 2015; Shimizu et al., 1998), the nuclear lamin B is disassembled through lamin B phosphorylation. To analyze the phosphorylation status of lamin B in NETotic neutrophils, we purified lamin B through immunoprecipitation and then analyzed the protein phosphorylation status with anti-p-Ser/Thr/Tyr antibody, since there is no relevant anti-phosphorylated lamin B (phospho-lamin B) antibody available for the direct detection of phospho-lamin B. Importantly, the increased phospho-lamin B was detected among total lamin B, purified through immunoprecipitation with anti-lamin B antibody, from NETotic neutrophils stimulated by PMA or PAF (Fig. 4F-G, Fig. S4A-B), as compared to those from control neutrophils.

Our results from immunoblot analysis with the isolated nuclei (Fig. 4A), or isolated extracellular neutrophil NETs (Fig. 4E), and confocal microscopy analysis of the nucleus of NETotic neutrophils (Fig. 3A, 4B-D), as well as the finding of lamin B phosphorylation among the purified total nuclear lamin B from NETotic neutrophils (Fig. 4F-G, Fig. S4A-B), indicate that nuclear accumulation of p-PKCα (Fig. 4A) may cause nuclear lamin B phosphorylation (Fig. 4F-G, Fig. S4A-B), and the phosphorylation-dependent lamin B disassembly that allows the intact full-length lamin B releasing with the neutrophil NETs to the extracellular space (Fig. 4E) from the ruptured nuclear envelope during NETosis.

### Decreasing lamin B phosphorylation by PKCα inhibition, or mutation of the consensus PKCα serine phosphorylation sites in lamin B, attenuates neutrophil NETosis

To explore the causal role of lamin B phosphorylation, we found that pre-treatment of human neutrophils with Go6976, a selective inhibitor of conventional PKC, attenuated PKCα phosphorylation in human neutrophils that were stimulated by PMA (Fig. 5A) or PAF (Fig. S5A). Importantly, inhibition of PKCα also decreased lamin B phosphorylation in immunoblot analysis with the purified nuclear lamin B immunoprecipitated from neutrophils that were treated by PMA (Fig. 5B) or PAF (Fig. S5B), without or with Go6976 pretreatment. Furthermore, pre-treatment of neutrophils with Go6976 decreased the nuclear accumulation of p-PKCα and prevented the nuclear envelope disintegration in PMA-stimulated neutrophils under confocal microscopy analysis (Fig. 5C). Consequently, inhibition of PKCα significantly attenuated NETosis of primary human pPMNs that were treated by PMA or PAF (Fig. S5C-D).

**Figure 5.**
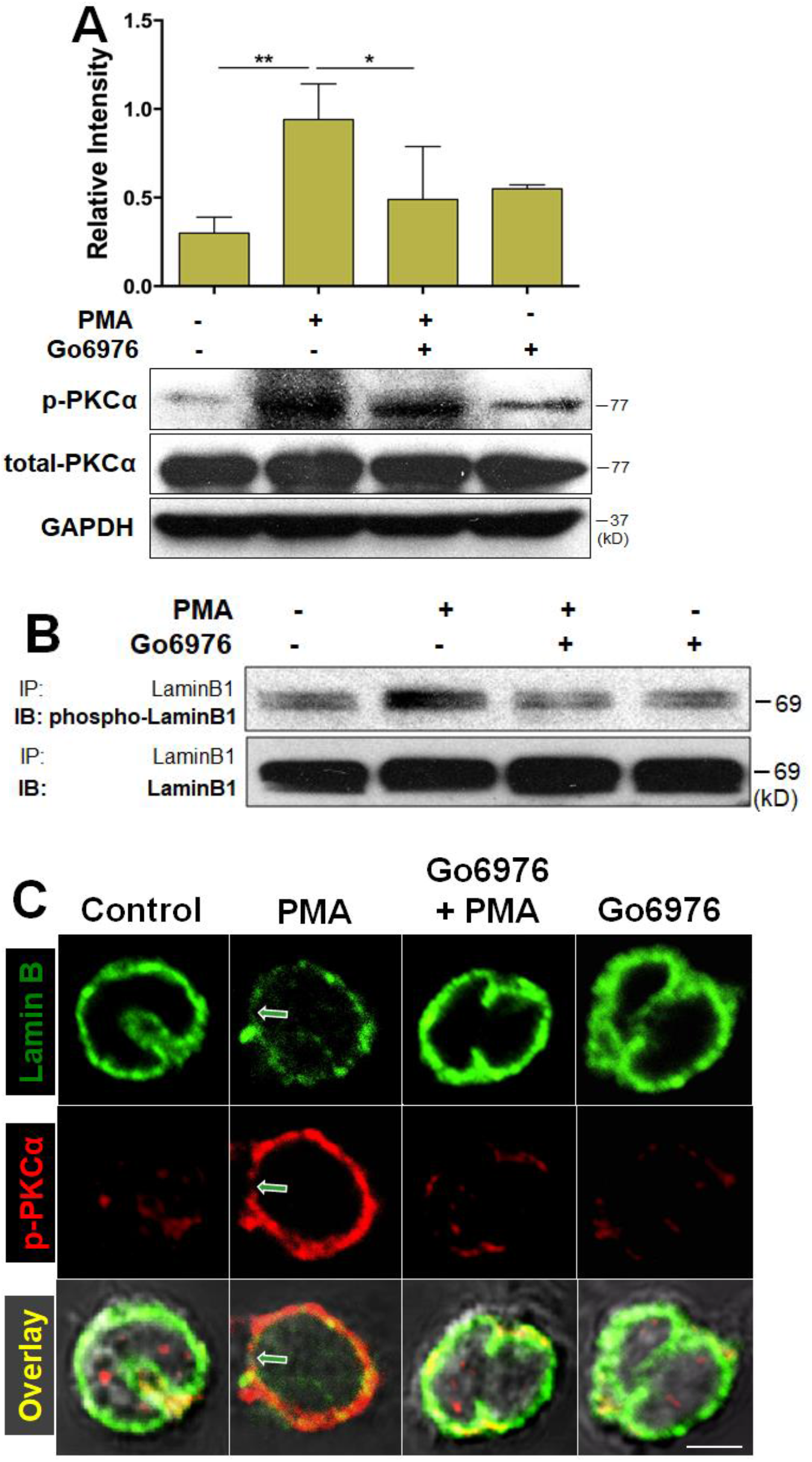
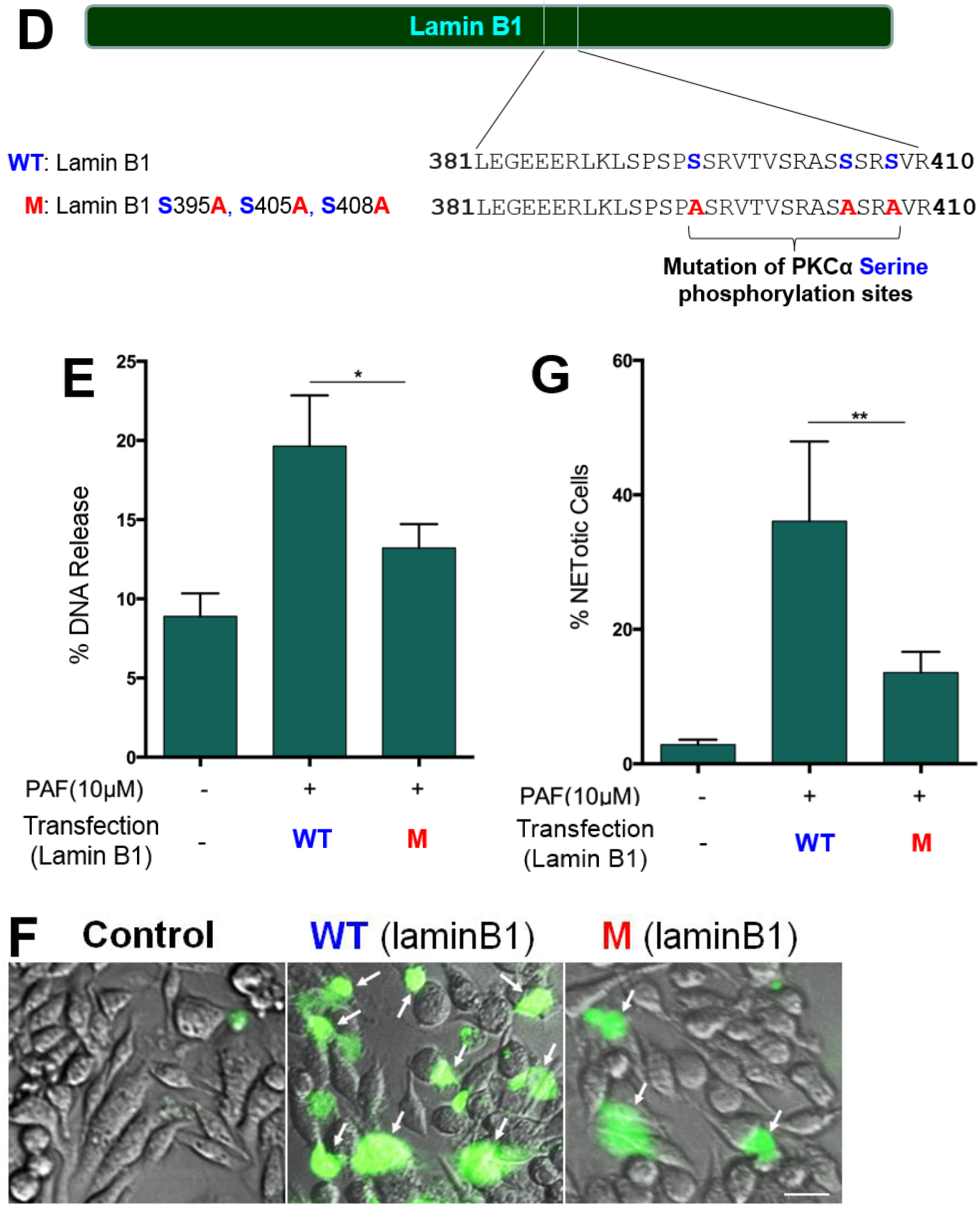
Inhibition of lamin B phosphorylation, or mutation of phosphorylation sites in lamin B, attenuates nuclear envelope rupture and neutrophil NET release. (**A**) Summary and representative immunoblots of total PKCα and p-PKCα in dPMNs that were pretreated without or with PKC inhibitor Go6976 for 1h, and then treated without or with PMA (A) for 3h. (**B**) Immunoblot (IB) detection of the phospho-lamin B and total lamin B with lamin B protein purified by immunoprecipitation (IP) with anti-lamin B from human dPMNs that were pretreated without or with PKC inhibitor Go6976 for 1h, and then treated by PMA (B) for 3h. (**C**) Confocal microscopy images of human dPMNs that were pretreated without or with PKCα Go6976 for 1h, and then treated without or with PMA for 3h, followed by staining of lamin B (primary anti-lamin B, and **FITC-**labeled secondary antibody), and phosphorylated PKCα (primary anti-human p-PKCα^S657^, and **PE**-labeled secondary antibody). The green/white arrows indicate the site of nuclear envelope rupture. Scale bars, 10 μm (C). (**D**) Schematic representation of lamin B1 domain structure and several mitotic serine phosphorylation sites (indicated by blue font) that exhibit PKC consensus motifs (Pearson and Kemp, 1991) and overview of lamin B1 reporters containing serine (S) to alanine (A) mutations at PKC consensus phosphorylation sites. (**E**) The endpoint analysis of NET-DNA release index of extracellular trap formation was detected for RAW264.7 cells that were transfected with plasmids of either wildtype lamin B (WT control), or mutants of PKCα-consensus-phosphorylation sites (S395A/S405A/S408A) of lamin B, and treated by PAF 10 μM in medium containing 1 μM Sytox Green dye with recording by a microplate reader at the 3h time point. RAW264.7 cells without transfection and without PAF treatment were served as basal control. The endpoint analysis of NET-DNA release index was reported in comparison to an assigned value of 100% for the total DNA released by RAW264.7 cells lysed by 0.5% (v/v) Triton-X-100. (**F,G**) The representative images (F) and immunofluorescent imaging quantification (G) of extracellular trap formation in the RAW264.7 cells that were transfected with plasmids of either wildtype lamin B (WT control), or mutants of PKCα-consensus-phosphorylation sites (S395A/S405A/S408A) of lamin B, and treated by 10 μM PAF for 3h. RAW264.7 cells without transfection and without PAF treatment were served as basal control. Then NETotic cells were stained by cell impermeable dye SYTOX Green (500 nM), and the total cells were detected by staining with cell-permeable dye SYTO Red (500 nM) and phase-contrast images. Images were taken by Olympus confocal microscopy, followed by automated quantification of NETs using ImageJ for quantification of % NETotic cells. Scale bars, 50 μm (C). Panels A,E,G are summary analyses of immunoblots that were calculated based on the arbitrary unit (A), or the NET-DNA release index calculated based on fluorescent readout (E), or percentage of NETotic cells by immunofluorescent imaging quantification using ImageJ (G), from more than 3 independent experiments. P*<0.05, P**<0.01, between groups as indicated. Comparisons amongst three or more groups were performed using ANOVA, followed by Student-Newman-Keuls test.

To further define the causal role of PKCα-mediated lamin B phosphorylation in NETosis, we made mutations at the PKCα-consensus serine phosphorylation sites (S395, S405, S408) of lamin B (Hocevar et al., 1993; Machowska et al., 2015; Mall et al., 2012) by replacement of serine (S) with alanine (A), which will prevent PKCα-mediated serine phosphorylation at lamin B. Mutants with single or multiple mutations (S395A, S395A/S405A, S395A/S405A/S408A) (Fig. 5D) were made. We found that extracellular trap formation was impaired in RAW264.7 cells that were transfected with plasmids containing mutations at all three (S395A/S405A/S408A), but not fewer (S395A, S395A/S405A, data not shown), PKCα-consensus serine phosphorylation sites of lamin B, as compared to those transfected with plasmids of WT lamin B, detected by the endpoint analysis of NET-DNA release index (Fig. 5E) and fluorescent imaging % NETotic cell analysis (Fig. 5F,G). In addition, pretreatment of RAW264.7 cells with PKCα inhibitor Go6976 attenuated PAF-induced extracellular trap formation (Fig. S5E). Furthermore, inhibition of PKCα by Go6976 decreased NET formation in neutrophils that were treated by other stimulus, cholesterol enrichment (Fig. S5F) (Liu et al., 2014). All of the above experiments indicate a causal link in which PKCα serves as a NETotic lamin kinase that mediates lamin B phosphorylation, following by the consequent nuclear envelope rupture, and release of extracellular trap in various cell types stimulated by different stimuli.

### Genetic deficiency of PKCα decreases lamin B phosphorylation and NETosis in primary mouse neutrophils from PKCα deficient mice

Next, we sought to further explore the causal effect of PKCα in lamin B phosphorylation and NETosis by using primary mouse peritoneal or bone marrow neutrophils from both heterozygous and homozygous PKCα deficient mice (Fig. 6A). Firstly, we found the decreased lamin B phosphorylation of the purified lamin B protein that was immunoprecipitated from the PKCα^-/-^ peritoneal mPMNs as compared to the WT mPMNs (Fig. 6B) without or with PAF treatment. These results further defined that PKCα serves as a NETotic lamin kinase and phosphorylates lamin B during NETosis induction. Most importantly, we found that both peritoneal (Fig. 6C-E) and bone marrow (Fig. S6A) mPMNs from either heterozygous (Fig. 6C) or homozygous (Fig. 6C-F) PKCα deficient mice had significantly impaired NETosis detected by conventional fluorometric NET quantification (Fig. 6C), the endpoint analysis of NET-DNA release index (Fig. 6D), or the immunofluorescent imaging quantification analysis (Fig. 6E-F). In addition, the protective effects of PKCα deficiency were dose-dependent as less neutrophil NETosis was seen in the neutrophils from homozygous PKCα deficient mice than those from heterozygous PKCα deficient mice (Fig. 6C, Fig. S6A). Therefore, our results from the genetic PKCα deficient mice provide definitive evidence that PKCα is important in NETosis, at least in part by serving as a NETotic lamin kinase through its regulatory role in phosphorylation and disassembly of lamin B, as well as the consequent nuclear envelope rupture in NETotic neutrophils.

**Figure 6.**
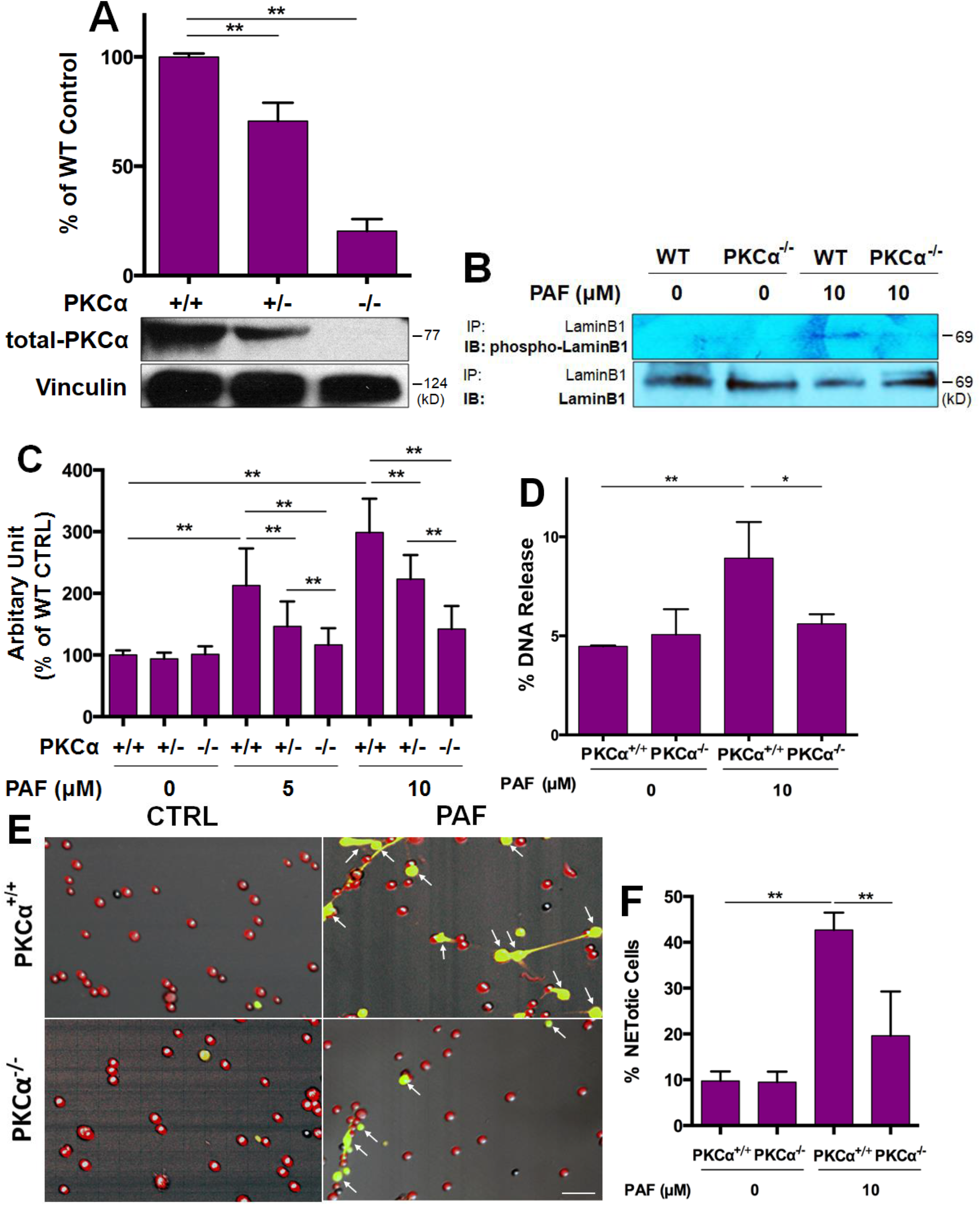
Genetic deficiency of PKCα attenuates lamin B phosphorylation and NETosis in primary neutrophils from PKCα knockout mice. (**A**) Summary and representative immunoblots of PKCα expressions in mouse bone marrow neutrophils from PKCα^-/-^, PKCα^+/-^, and their littermate WT (PKCα^+/+^) mice. Vinculin served as loading control. (**B**) Immunoblot (IB) detection of the phospho-lamin B and total lamin B with lamin B protein purified by immunoprecipitation (IP) with anti-lamin B from mouse peritoneal mPMNs from WT vs PKCα^-/-^ mice, stimulated without or with PAF for 3h. (**C**) Summary analyses of NETosis in peritoneal mPMNs, from WT, PKCα^+/-^, or PKCα^-/-^ mice, which were treated without or with either 5 µM or 10 µM PAF for 3h following by 2% PFA fixation and fluorometric microplate analysis. (**D**) The endpoint analysis of NET-DNA release index was detected by coincubation of primary mouse peritoneal mPMNs from WT and PKCα^-/-^ mice without (control) or with 10 µM PAF in medium containing 1 μM Sytox Green dye with recording by a microplate reader at the end of the 3h time point. The NET-DNA release index was reported in comparison to an assigned value of 100% for the total DNA released by neutrophils lysed by 0.5% (v/v) Triton-X-100. (**E**) Representative and (**F**) summary analysis of NETosis of mouse peritoneal mPMNs from WT vs PKCα^-/-^ mice stimulated without or with 10 µM PAF for 3h, and stained with both cell permeable Syto Red dye and cell impermeable Sytox Green dye, without fixation. Images were taken by Olympus confocal microscopy, followed by automated quantification of NETs on 5-6 non-overlapping area per well using ImageJ for calculation of % NETotic cells. (D) Scale bars, 50 μm. Panels A,C,D,F were summary analyses that were calculated based on the arbitrary unit (A) or arbitrary fluorescent readout (C), endpoint analysis of NET-DNA release index (D), or % NETotic cells by image analysis using ImageJ (F) from more than 3 independent experiments as compared to their WT or untreated controls, P*<0.05, P**<0.01 between groups as indicated. Comparisons amongst three or more groups were performed using ANOVA, followed by Student-Newman-Keuls test.

### Strengthening nuclear envelope integrity by lamin B overexpression attenuates neutrophil NET release in vivo and alleviates exhibition of NET-associated proinflammatory cytokines in UVB-irradiated skin of lamin B transgenic mice

Here, we first showed the involvement of neutrophil NETosis in inflammatory responses of UVB-induced skin inflammation in WT mice after UVB irradiation (Fig. 7A-B). As we found a crucial role for lamin B integrity in neutrophil NETosis *in vitro* (Fig. 1E-G), here we sought to verify the above findings with *in vivo* experiments. We found that UVB exposure induced inflammatory responses in mouse skin with increased infiltration of neutrophils (Fig. 7A). Many of those infiltrated neutrophils became NETotic in the dermis of WT littermates after UVB irradiation (Fig. 7B-C, Fig. S7A). Confocal microscopy analyses revealed the extensive extrusion of decondensed chromatin to the extracellular space of the NETotic neutrophils in the dermis of the UVB-irradiated WT littermates (Fig. 7C). In contrast, we observed fewer neutrophils, particularly the non-netting swelling neutrophils that enclosed citrullinated chromatin (anti-cit H3 staining) inside, with no chromatin extrusion, of the cells in the UVB-irradiated skin of the Lmnb1^TG^ mice (Fig. 7D). These *in vivo* experimental results confirm the findings from the *in vitro* study (Fig. 1E-G), in which strengthening nuclear envelope integrity by lamin B overexpression attenuated NET release in neutrophils. Furthermore, the NETotic neutrophils exhibited proinflammatory cytokines, IL-17A and TNFα, in the UVB-irradiated skin of WT littermates (Fig. 7E-F, Fig. S7B-C). Most importantly, the attenuated neutrophil infiltration, reduced neutrophil NETosis, and alleviated exhibition of NET-associated proinflammatory cytokines, were demonstrated in the UVB-irradiated skin of the Lmnb1^TG^ mice (Fig. 7C-F, Fig. S7B-C), as compared to those in their WT littermates. It has been known that IL-17A and TNFα can synergistically work together to sustain neutrophil recruitment during inflammatory responses (Griffin et al., 2012; Liu et al., 2011). Therefore, the reduced release of neutrophil NETs with decreased extracellular display of IL-17A and TNFα may explain the attenuated neutrophil recruitment/infiltration to the UVB-irradiated skin in LmnB1^TG^ mice as compared to their WT littermates.

**Figure 7.**
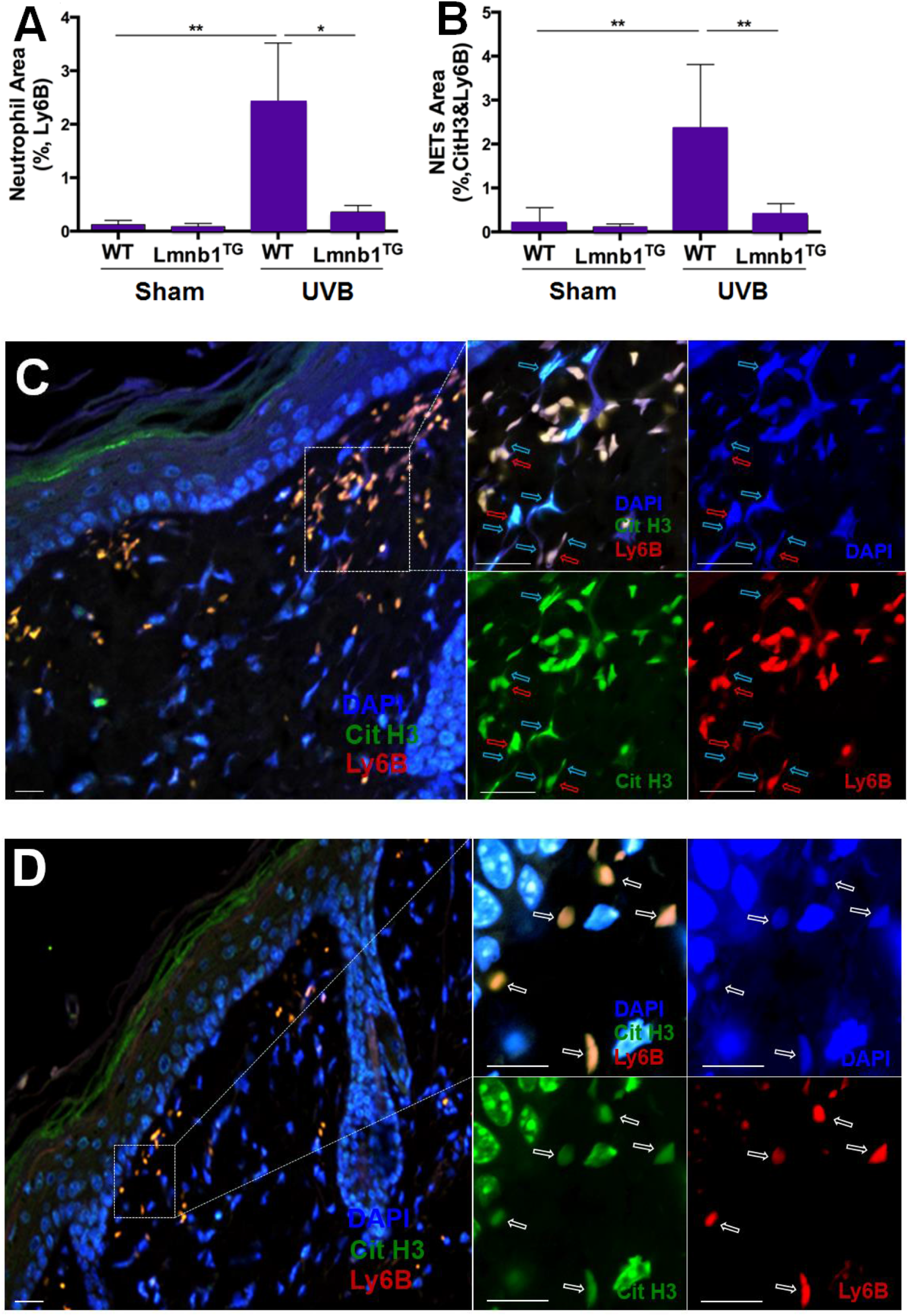
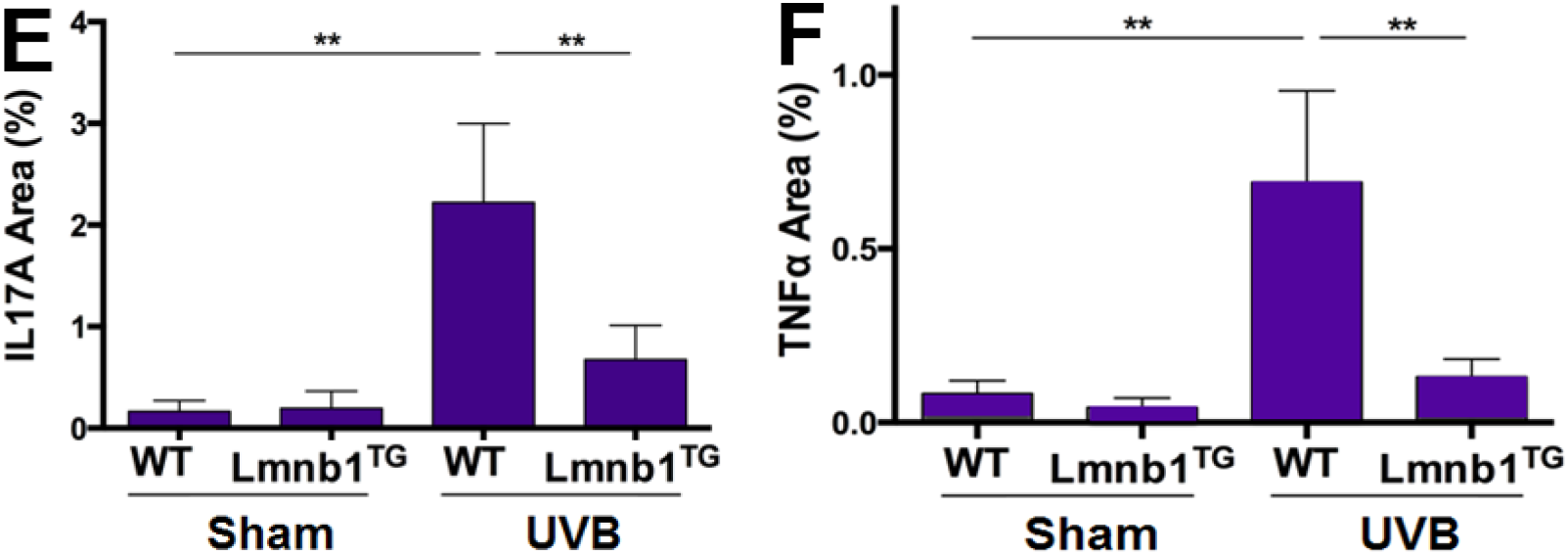
Strengthening nuclear envelope integrity by lamin B overexpression decreases neutrophil NET release in vivo and alleviates NET-associated proinflammatory cytokine accumulation in UVB irradiated skin of the lamin B transgenic mice. (**A**) Summary analysis of neutrophil staining area in the skin tissue by detection of the area of neutrophils that were stained by rat anti-mouse Ly6B, and Alexa Fluor 647 labeled goat anti-rat IgG secondary Ab. (**B**) Summary analysis of NETotic neutrophils. and (**C**,**D**) representative confocal fluorescent microscopy images of NETotic neutrophils. in the skin tissue of WT (C) vs Lmnb1^TG^ (D) mice with UVB exposure, in which DNA was stained by **DAPI**, citrullinated histone H3 was probed by rabbit anti-mouse citrullinated Histone H3, following stained by **Alexa Fluor-488** labeled donkey anti-rabbit secondary antibody, while neutrophil surface marker Ly6B was probed by Rat anti-mouse Ly6B Ab following stained by **Alexa Fluor-647** conjugated goat anti-rat secondary antibody. The images of different staining and their overlays are shown (C,D). The parental neutrophils (stained by neutrophil membrane surface marker Ly6B) and their released extracellular NET structures (co-stained by anti-cit H3 and DAPI) are indicated by red arrows and light blue arrows respectively. The white arrows indicated the non-netting neutrophils. Scale bar, 50 µm. (**E**,**F**) Summary analysis for staining area of NETotic neutrophils with their exhibition of IL-17A (E) or TNFα (F), in the skin tissue by co-staining with anti-Ly6B, or the IL-17A (E) or TNFα (F) fluorescent staining area that were stained with rabbit anti-mouse IL-17A, followed by Alexa Fluor 488 labeled donkey anti-rabbit IgG Ab, or stained by FITC conjugated rat anti-mouse TNFα Ab. Panels A,B,E,F are summary analyses that were calculated based on the arbitrary fluorescent unit (A,B,E,F) from more than 3 independent experiments of different groups as indicated. P*<0.05, P**<0.01 between groups are indicated. Comparisons amongst three or more groups were performed using ANOVA, followed by Student-Newman-Keuls test.

The results from all of the above *in vitro* and *in vivo* studies have verified our hypothesis that lamin B is crucial to nuclear envelope integrity and the nuclear envelope rupture during neutrophil NET release. Therefore, strengthening nuclear envelope integrity by lamin B overexpression attenuated neutrophil NET release *in vitro* and *in vivo* in lamin B transgenic mice.

## DISCUSSION

Externalization of decondensed chromatin from ruptured nuclear envelope forms the backbone of neutrophil NETs. Lamin B filamentous meshwork is a structural component of the nuclear envelope and anchored to the inner nuclear membrane by lamin B receptor (LBR) (Figure 1C) (Olins et al., 2010). Given the nature of their structural arrangement, lamin B filaments are assembled laterally to form a highly organized meshwork underneath the inner nuclear membrane, the two are vertically associated by the integral membrane protein LBR (Figure 1C) (Goldberg et al., 2008; Vergnes et al., 2004). Therefore, the lateral dimensional lamin B meshworks might be more important in maintaining nuclear envelope integrity and enclosing nuclear chromatin within the nucleus than do the LBR. The role of lamin B in neutrophil biology and its involvement in NETotic nuclear envelope rupture has not been investigated, although LBR has been used as a marker of the nuclear envelope in neutrophil biology, including NETosis, studies (Fuchs et al., 2007; Singh et al., 2016). In the current study, we verified the existence of lamin B both in human and in mouse neutrophils. Nuclear lamin B is involved in NETotic nuclear envelope rupture and the disassembled nuclear lamin B is decorated on the surface of the externalized nuclear chromatin during the release of neutrophil NET from the ruptured nuclear envelope. To test the effects of levels of nuclear lamin B on nuclear envelope integrity, we found that overexpression of lamin B strengthened the nuclear envelope integrity and blunts the NET release *in vitro*. In contrast, the reduced amount of mature nuclear lamin B by inhibition of protein farnesylation with the farnesyltransferase inhibitor (FTI) (Adam et al., 2013) can enhance the release of extracellular traps. Taken together, the levels of nuclear lamin B expression affect extracellular trap formation in a negative dose-dependent manner. Therefore, our findings provide definitive evidence for the importance of lamin B in nuclear envelope integrity and NET release in neutrophils. In line with our findings, a recent work by Zychlinsky and colleagues reported that lamin A/C is involved in neutrophil NETosis (Amulic et al., 2017).

In the literature, nuclear lamin B is either cleaved or disassembled during nuclear envelope rupture in different cellular events (Collas, 1998; Hocevar et al., 1993; Slee et al., 2001). According to the lytic cell death mechanism of NETosis described before (Fuchs et al., 2007; Papayannopoulos et al., 2010; Yipp and Kubes, 2013), one may expect to see the destructive proteolytic cleavage of lamin B for the ruptured nuclear envelope in NETotic neutrophils. Unexpectedly, we found that lamin B remained as an intact full-length molecule not only in the whole-cell lysates of the NETotic neutrophils, but also in the externalized NET structure isolated from the conditioned medium of NETotic neutrophils, indicating that this nuclear structural filamentous protein was not destructively cleaved during the NETotic nuclear envelope rupture. In contrast, lamin B in the apoptotic neutrophils was fragmented, which is known to be cleaved by proteolytically activate caspase-3 (Cross et al., 2000; Kivinen et al., 2005). While pro-caspase-3 remained as an inactive molecule throughout the period of neutrophil NETosis induction in our study, further explaining the nature of the uncleaved full-length lamin B in the ruptured nuclear envelope in NETosis. Our results are in line with the previous report in which NETosis is a caspase-independent process (Remijsen et al., 2011). All of the above results suggest that nuclear lamin B was not destructively cleaved during NETotic nuclear envelope rupture. Therefore, our results are in line with several other studies (Amulic et al., 2017; Neubert et al., 2018) and suggest that the rupture of the nuclear envelope appears to be a distinct process from the previously described lysis or dissolution of the nuclear envelope (Fuchs et al., 2007; Papayannopoulos et al., 2010; Yipp and Kubes, 2013).

It has long been known that polymerization or assembly of the monomeric lamin proteins (lamin A/C, B) into fibers and complex lamina meshworks strengthen nuclear envelope integrity (Aebi et al., 1986; Holaska et al., 2002). Given the nature that the intact full-length lamin B and inactive pro-caspase 3 were found in the NETotic neutrophils in our study, we proposed the potential disassembly of lamin B might be responsible for the nuclear envelope rupture during NETosis. To explore the potential mechanisms that regulate nuclear lamin B disassembly in NETosis, we found that cytoplasmic PKCα was translocated to the nucleus where it became phosphorylated and activated. It is known that nuclear translocation and activation of PKCα is stimulated by nuclear diacylglycerol, derived from nuclear membrane lipids (Irvine, 2000; Martelli et al., 2006; Neri et al., 1998), and phosphorylation of PKCα at Ser-657 can serve as a marker of PKCα activation (Bornancin and Parker, 1997; Gysin and Imber, 1996). Therefore, our results of accumulation of p-PKCα^Ser657^ in the nucleus may represent the activation of PKCα that is responsible for lamin B phosphorylation and phosphorylation-dependent disassembly (Edens et al., 2017; Hocevar et al., 1993; Hornbeck et al., 1988), as well as the consequent NETotic nuclear envelope rupture. The experimental evidence in the current study, that support this notion, includes: 1) the concomitant detection of p-PKCα and full-length lamin B in the nuclear fraction of NETotic neutrophils and the isolated extracellular NETs, 2) the accumulation of p-PKCα at the site of nuclear envelope rupture in NETotic neutrophils, 3) the demonstration of phosphorylated lamin B that was purified from NETotic neutrophils with immunoprecipitation. In the current study, co-existence of p-PKCα and lamin B in the nucleus indicate their close interaction that may cause lamin B phosphorylation since lamin B is the major nuclear substrate for PKC-mediated phosphorylation (Collas et al., 1997). Studies from others (Collas et al., 1997; Fields et al., 1988; Hornbeck et al., 1988), and from ourselves (Mall et al., 2012) have demonstrated that phosphorylation of lamin B results in disassembly of this filament protein, and consequent nuclear envelope breakdown. Therefore, our findings in the current study indicate an important role of PKCα in lamin B phosphorylation and disassembly, as well as the consequent nuclear envelope rupture and neutrophil NET release.

To determine the causal role of PKCα, firstly we found that pharmacological inhibition of PKCα attenuated PKCα phosphorylation, with the reduced appearance of p-PKCα in the nucleus and decreased disintegration of nuclear envelope under the confocal microscopy, as well as the decreased lamin B phosphorylation, and the reduced neutrophil NETosis. Secondly, and most importantly, mutation of three, but not fewer (data not shown), PKCα-consensus serine phosphorylation sites of lamin B impaired the release of extracellular traps, indicating the causal role of PKCα-mediated lamin B phosphorylation in lamin B disassembly and nuclear envelope rupture during chromatin release and extracellular trap formation. Furthermore, genetic deficiency of PKCα significantly inhibited phosphorylation of nuclear lamin B purified by immunoprecipitation, and NETosis in neutrophils from PKCα KO mice. Taken together, the above results with different approaches provide evidence for the causal role of PKCα in nuclear lamin B phosphorylation and neutrophil NETosis. Previous studies have documented that PKC may be important in neutrophil NETosis, probably through inhibition of the PKC-Raf-ERK/MAPK signaling pathway (Hakkim et al., 2011), or NOX-mediated ROS generation (Fuchs et al., 2007; Hakkim et al., 2011; Papayannopoulos et al., 2010). In the current study, we identified a causal role of PKCα in NETosis and elucidated a novel mechanism by which PKCα serves as a lamin kinase that mediates nuclear lamin B phosphorylation and disassembly, and the consequent NETotic nuclear envelope rupture for externalization of decondensed nuclear chromatin and neutrophil NET formation.

The externalization of nuclear chromatin forms the backbone of neutrophil NETs (Brinkmann et al., 2004; Papayannopoulos et al., 2010), which is decorated with other cytosolic contents, granule proteins, and even mitochondrial DNA during the process of nuclear chromatin extrusion and neutrophil NET formation (Lood et al., 2016). Based on these facts, nuclear chromatin decondensation and nuclear envelope rupture are required for neutrophil NET formation. It is already known that nuclear chromatin is either decondensed by PAD4-catalyzed histone citrullination (Neeli et al., 2008; Wang et al., 2009), or degraded by neutrophil elastase-mediated histone degradation (Dhaenens et al., 2014; Fuchs et al., 2007; Papayannopoulos et al., 2010). As a pre-requisite for extrusion of the decondensed chromatin, the nuclear envelope rupture is required, but achieved via an unknown mechanism (Papayannopoulos, 2017). With genetic manipulation and pharmacological approaches, the current study identified that nuclear lamin B is crucial to the integrity of nuclear envelope. PKCα serves as lamin kinase that mediates lamin B phosphorylation and disassembly, and the consequent nuclear envelope rupture and neutrophil NET release. In agreement with our novel findings, a recent work from Zychlinsky and colleagues reported that cyclin dependent kinases (CDK) 4/6 might be important in the regulation of neutrophil NETosis through phosphorylation of lamin A/C (Amulic et al., 2017). In addition, Neubert et al reported that nuclear chromatin swelling provides physical driving force that causes the nuclear envelope rupture and release of neutrophil NETs (Neubert et al., 2018). However, all of these latest novel findings from our and other groups (Amulic et al., 2017; Neubert et al., 2018) do not support the lytic hypothesis of nuclear envelope rupture in NETosis, at least in the early stage of this programmed cell death (Neubert et al., 2018).

Neutrophils are known to be the first group of cells to be recruited to the site of acute inflammation (Kruger et al., 2015), including skin inflammation induced by UVB irradiation (Hawk et al., 1988). In the current study, we found NETotic neutrophils with extensive extrusion of citrullinated chromatin, and their exhibition of proinflammatory cytokines, IL-17A and TNFα by the extracellular NETs in the dermis of UVB-irradiated WT mice. It is known that UVB irradiation can induce IL-17A (Li et al., 2015; MacLeod et al., 2014) and TNFα (Bashir et al., 2009; Sharma et al., 2011), and neutrophils can release IL-17A (Lin et al., 2011; Tecchio et al., 2014) and TNFα (Giambelluca et al., 2014; Tecchio et al., 2014). Neutrophil NET-associated pro-inflammatory cytokines may survive longer and distribute in a relatively higher concentration within the microenvironment around the NETs in the skin than do their soluble forms that may be quickly diluted in circulating blood. Additionally, co-exhibition of IL-17A and TNFα by neutrophil NETs may propagate inflammatory responses as the two are known to have synergistic proinflammatory effects in immune cell recruitment (Griffin et al., 2012; Liu et al., 2011), or production of other cytokines (Ruddy et al., 2004). Importantly, strengthening nuclear envelope integrity by lamin B overexpression attenuated neutrophil NET release *in vivo*, in which the infiltrated neutrophils were non-netting swelling and without extensive chromatin release in the skin of UVB-irradiated lamin B transgenic mice. The reduced NET release in skin is accompanied by decreased extracellular exhibition of proinflammatory cytokines, IL-17A and TNFα, by the non-netting neutrophils, which may explain the attenuated inflammatory cell infiltration in skin of lamin B transgenic mice following UVB irradiation. In the current study, we focused on the cellular mechanisms that lamin B phosphorylation is crucial to nuclear envelope rupture and NET release. Although the *in vivo* experiments in the current study have shown the reduced NET-associated inflammatory cytokine exhibition by the attenuated neutrophil NETosis in LmnB1^TG^ mice, additional investigations are needed in the future to further elucidate the mechanisms of the reduced skin inflammation regarding the relevant changes, other than neutrophils, i.e. keratinocytes and other inflammatory cells, in LmnB1^TG^ mice following UVB-irradiation.

In conclusion, our novel findings identified that 1) lamin B is crucial to the nuclear envelope integrity in neutrophils, 2) PKCα serves as NETotic lamin kinase that mediates nuclear lamin B phosphorylation and disassembly, following by the consequent nuclear envelope rupture and neutrophil NET release, 3) strengthening nuclear envelope integrity by lamin B overexpression attenuates neutrophil NETosis *in vitro* and *in vivo*, and alleviates NET-associated proinflammatory cytokine exhibition in skin of the UVB-irradiated lamin B transgenic mice. Our novel findings advance our understanding of NETosis process and highlight a cellular mechanism in which PKCα-mediated lamin B phosphorylation drives nuclear envelope rupture for NET release in NETotic neutrophils. Therefore, our work combining few other recent works (Amulic et al., 2017; Neubert et al., 2018) provide new insights into a potential therapeutic strategy for the neutrophil NET associated human diseases.

## MATERIALS AND METHODS

### Mouse

Lamin B1 transgenic C57BL/6J-Tg (Lmnb1^TG^) 1Yfu/J mice (JAX stock #023083) (Heng et al., 2013), PKCα deficient mice (B6;129-Prkca ^tm1Jmk^/J mice, JAX stock #009068) (Braz et al., 2004), and their corresponding littermate controls, were housed in a pathogen-free environment, and given food and water *ad libitum*. All the animal experiments were approved by the Animal Care and Use Committee of Philadelphia VA Medical Center.

### Cell Culture and Treatment

Primary human polymorphic nuclear granulocytes (pPMNs) or neutrophils were isolated by *dextran* (250,000) sedimentation of red blood cells (RBC) followed by Ficoll-Paque PLUS H-1077 (GE Healthcare) gradient separation as described before with modification (Denny et al., 2010). Human HL-60 cell line was differentiated into neutrophil-like or polymorphic nuclear granulocytes (dPMNs) (Birnie, 1988) with 1.2% DMSO for 5-6 days as described in our published work (Folkesson et al., 2015). Primary mouse peritoneal and bone marrow neutrophils (mPMNs) were isolated according to the published protocols (Braz et al., 2004). PKCα deficient mPMNs were isolated from heterozygous (PKCα^+/-^) or homozygous (PKCα^-/-^) PKCα deficient mice (B6;129-Prkca ^tm1Jmk^/J mice, JAX stock #009068) (Braz et al., 2004), while mPMNs with lamin B1 overexpression (Lmnb1^TG^) were isolated from lamin B1 transgenic C57BL/6J-Tg(LMNB1) 1Yfu/J mice (JAX stock #023083) (Heng et al., 2013). Neutrophils from their littermate wildtype (WT) control mice for both PKCα deficient and lamin B1 transgenic strains served as controls. Mouse peritoneal mPMN isolation was conducted with 3% thioglycollate broth (TGB) medium induction, and then isolated with neutrophil isolation kit (Miltenyl Biotec) according to manufacturer’s instruction. Mouse bone marrow mPMNs were isolated from the *femur* and *tibia* bone marrow with a neutrophil isolation kit (Miltenyl Biotec). RAW264.7 cells (ATCC) were cultured in DMEM medium supplemented with 10% FBS, and were also used in certain experiments. All of the above pPMNs, dPMNs, mPMNs, or RAW264.7 were treated with, either the most commonly used NETosis inducer PMA (50 or 100 nM), or the naturally occurring lipid mediator PAF (5 or 10 µM), for 0-3h without or with 1h pretreatment with an inhibitor of conventional PKC Go6976, following by detection of NETosis, or lysis for immunoblot analysis. In addition, cholesterol loading of pPMNs was used as other stimulus for induction of NETosis, and conducted by coculture of the cells with water-soluble cholesterol (100 µg/ml) complex with methyl-β-cyclodextrin for 6h (Liu et al., 2007). Since lamin B maturation is regulated by farnesylation (Adam et al., 2013), the reduced mature lamin B1 expression in RAW264.7 cells was achieved by treatment without (0) or with 2 or 10 μM farnesyltransferase inhibitor (FTI) L-744,832 for 48h.

### Neutrophil NET Formation Assessment and Quantification

All of the above pPMNs, dPMNs, or mPMNs were treated with, either the most commonly used NETosis inducer PMA (50 or 100 nM), or the naturally occurring lipid mediator PAF (5 or 10 µM), for 0-3h. Three currently validated methodologies were used for NETosis assessment and quantification in the current study. **1**) In the conventional fluorometric NET quantification (Chicca et al., 2018; Sollberger et al., 2016), the above treated or un-treated cells were fixed with 2% PFA after the stimulation, and stained with 1 µM SYTOX Green, following by fluorescent readout with a microplate reader at 504/523 nm as described before with modification (Chicca et al., 2018; Pang et al., 2013; Sollberger et al., 2016). NET formation in each experiment was also confirmed by fluorescent microscopy. **2**) For the NET-DNA release index under different conditions (Douda et al., 2015; Khan et al., 2018), the cells without (control) or with PMA or PAF treatment for 3h in medium containing 1 μM Sytox Green dye at 48-well plates. The experiments were recorded by a microplate reader (504/523 nm) either at the end of the 3h treatment for endpoint analysis, or every 20 min for up to 3h for kinetic analysis. The NET-DNA release index was reported in comparison to an assigned value of 100% for the total DNA released from cells lysed by 0.5% (v/v) Triton-X-100 as described before with modification (Douda et al., 2015; Khan et al., 2018). To calculate the kinetic NET-DNA release index in each condition, the fluorescence intensity at time 0 min was subtracted from the fluorescence at each time point and was then divided by the fluorescence values of cell lysed with 0.5% (v/v) Triton X-100. All the experimental values were standardized to total DNA released by cell lysis. Each condition was tested with a technical triplicate. **3**) For the immunofluorescent imaging quantification analysis of NETosis (Martinod et al., 2017; Sollberger et al., 2016), the equal number of cells were treated by stimuli for 3h. Then cell impermeable DNA dye SYTOX Green (500 nM) was used to detect NETotic cells, while the total number of cells were determined by staining with cell-permeable DNA dye SYTO Red (500 nM) and phase-contrast imaging. All images were acquired with Fluoview 4.2 software by Olympus confocal microscopy, followed by automated quantification of NETs on 5-6 non-overlapping area per well using ImageJ for quantification of % NETotic cells.

### Neutrophil NET Isolation

Neutrophil NETs isolation was processed according to the published literature (Najmeh et al., 2015) with modification. Briefly, the conditioned medium of NETotic neutrophils were gently aspirated and discarded after NETs generation. Cold PBS without Ca^2+^ and Mg^2+^ was used to collect NETotic neutrophils solution. All cells were removed by centrifuge at 450 x g for 10 min at 4 °C; the cell-free NET-rich supernatant were collected and followed by centrifuge at 18,000 x g for 10 min at 4 °C. The isolated NETs lysed by RIPA lysis buffer containing phosphatase inhibitor cocktail and protease inhibitor cocktail (Roche) and the NET lysates were used for immunoblot assay.

### DNA Constructs and Cell Transfection

The pEGFP-C1-lamin B1 was used to generate phosphorylation site mutants. All plasmids for PKCα-consensus phosphorylation site mutant isoforms with single or multiple mutations (pEGFP-C1-lamin B1 S395A, pEGFP-C1-lamin B1 S395A/S405A, pEGFP-C1-lamin B1 S395A/S405A/S408A) were generated as described before (Mall et al., 2012). RAW264.7 cells (ATCC) were cultured in DMEM medium supplemented with 10% FBS, and were transfected with TransIT-Jurket (Mirus) according to the manufacturer’s instructions. Twenty-four hours after transfection, cells were treated by 10 μM PAF for 3 hours and evaluated for induction of extracellular traps, which were then detected as described above.

### Confocal Fluorescent Microscopy Analysis

Neutrophils and NETs were fixed with 100% methanol, then stained with primary anti-total PKCα, or anti-phospho-PKCα^S657^, or anti-lamin B Abs, followed by their matched secondary Abs with FITC or PE conjugation. DNA was stained with DAPI. For measurement of nuclear envelope continuity, the circumferences/perimeters of the cell nucleus, and the lengths of the discontinuity of ruptured nuclear envelope (lamin B staining), of the cells from confocal microscopy images were measured with ImageJ software (Evans et al., 2009) with modification. Then, the percentages of the discontinuity were calculated according to the above measurements. Confocal fluorescent images were analyzed with an Olympus Fluoview 1000 confocal microscope (Li et al., 2013; Liu et al., 2012).

### Nuclear Protein Extraction

To further study the nuclear accumulation of PKCα and its association with nuclear lamin B in neutrophils, the nuclei were extracted using hypotonic buffer according to the published protocol (Shaiken and Opekun, 2014) with modification. In brief, the collected neutrophils were re-suspended in ice-cold buffer (pH 7.4) containing 10 mM Tris-HCl, 10mM NaCl, and 5mM *magnesium* acetate for 30 minutes. NP-40 (0.3%) detergent was then added and homogenized. The mixture homogenates were centrifuged at 1,200 g for 10 minutes at 4℃, to collect sedimented nuclei; these were then resuspended in high volumes of buffer containing 880 mM sucrose and 5 mM magnesium acetate. The suspension was then centrifuged at 2,000 g for 20 minutes at 4℃ and nuclei were resuspended in buffer containing 340 mM sucrose and 5 mM magnesium acetate. The recovered nuclei were dissolved in 8 M urea, followed by sonication, and centrifugation by 10,000 g for 10 minutes at 4℃.

### Immunoprecipitation (IP)

Immunoprecipation was carried out with direct magnetic IP kit (Pierce) by coupling lamin B antibody (66095-1-Ig, Proteintech) to N-hydroxysuccinimide (NHS)-activated magnetic beads, followed by incubating the lysate with the coupled magnetic beads at 4°C overnight. The immunoprecipitates were then washed twice with cold washing buffer and once with cold ultra-pure water, then bound lamin B was eluted for immunoblot detection of phosphorylated and total lamin B, with anti-p-Ser/Thr/Tyr Ab and anti-lamin B Ab respectively, in neutrophils that were treated without or with PMA or PAF for 3h.

### Immunoblot Analysis

Immunoblots were conducted according to our published protocol (Li et al., 2013). Briefly, neutrophils that were treated or not were harvested and washed in ice-cold PBS, then lysed with RIPA lysis buffer containing phosphatase inhibitor cocktail and protease inhibitor cocktail (Roche). Equal amounts of protein were loaded on SDS-PAGE gel and electrophoresed, and then transferred to PVDF membranes. After membrane blocking, they were probed with primary antibodies and matched HRP-conjugated secondary antibodies, and then detected using an enhanced chemiluminescent detection kit (Millipore). In some of our immune-blot results, full-length lanes were shown to display the status of lamin B that was either intact or fragmented. Relative densitometric values of protein bands were quantified by the ImageJ 1.51f software (NIH, USA).

### Exposure of Mice to Ultraviolet B (UVB)

For the experiments with UVB exposure, the dorsal skin was exposed to UVB with two FS40T12-UVB bulbs (LIGHT SOURCES) according to our published protocol (Sharma et al., 2011) with modification. The lamin B1 transgenic Lmnb1^TG^ mice and their WT littermates in 8 weeks of age were received without/with 150 mJ/cm^2^ for 5 consecutive days under anesthetization by peritoneal injected Ketamine/Xylazine. Animals were sacrificed 24 hours after the last exposure. The whole dorsal skin samples, including epidermis, dermis, and subcutaneous fat were collected. Tissue sample sectioning was performed by the Penn Skin Biology and Diseases Resource-based Center.

### Fluorescent Immunohistochemistry

Immunohistochemistry staining was conducted with our previous established protocol with modification (Williams et al., 2005). Paraffin-embedded skin tissue samples were cut in to 5µm sections and placed on glass slides. Slides were placed in a 60°C oven overnight, deparaffinized with Citrisolv® (Fisher Scientific Waltham, MA) and rehydrated with serial ethanol dilutions. Samples underwent antigen retrieval using Target Retrieval Solutions (DAKO, Glostrup, Denmark), Peroxidase was deactivated using 0.3% H_2_O_2_ dissolved in deionized H_2_O for five minutes. Sections were blocked using DAKO protein block (Carpintera, CA). Samples were incubated with primary antibodies overnight at 4 °C. Primary antibodies used for staining included Ly6B (Rat anti-neutrophil [7/4], 1:500, Abcam, ab53457), Citrullinated histone H3 (citrulline R2+R8+R17) (Rabbit anti-citrullinated Histone H3, 1:300, Abcam, ab5103), IL-17A (Rabbit anti-IL-17A, 1:100, Abcam, ab79056). Goat anti-rat IgG secondary antibody conjugated with Alexa Fluor ®647 (1:500, Invitrogen), and donkey anti-rabbit IgG secondary antibody conjugated with Alexa Fluor ®488 (1:500, Invitrogen) were used to detect primary antibodies. FITC anti-mouse TNFα antibody (Rat anti-TNFα, 1:100, BioLegend, 506304), was also used for immunofluorescent staining. Isotype-matched IgG was used instead of primary antibody as a negative control of the staining. Counterstaining with DAPI was performed and slides were mounted with SlowFade Gold Antifade Mountant (Invitrogen). Images were obtained using the Nikon Eclipse fluorescent microscope, the fluorescent staining area percentage were analyzed by Nikon NIS-Elements Advanced Research software, according to previous publication (Brinkmann et al., 2016; Savchenko et al., 2014).

## QUANTIFICATION AND STATISTICAL ANALYSIS

Prism 6 (GraphPad Software) was used to perform statistical analysis. Normally distributed data are shown as means ± standard deviations. Comparisons between two groups were conducted with student T test. Comparisons amongst three or more groups were performed using ANOVA, followed by Student-Newman-Keuls test. Statistical significance was considered at a level of *P*-value < 0.05.

## SUPPLEMENTAL INFORMATION

Supplemental information includes seven figures can be found with this article online.

## AUTHOR CONTRIBUTIONS

M.L.L. conceived and designed the study. Y.L., M.L.L. performed the laboratory experiments and analyzed the experimental data. M.M. made the plasmids for lamin B mutation. M.L.L., Y.L., V.P.W., wrote/drafted and finalized the paper. All authors read and approved the final manuscript.

## ACKNOWLEDGEMENTS

The authors would like to acknowledge Penn Skin Biology and Diseases Resource-based Center for their skin histology service. The authors would thank Debra A. Pawlowski (animal facility, Philadelphia VA Medical Center) for her kindly help with our animal experiments. The authors would also thank Dr. Iain W. Mattaj for his support on the lamin B mutation work. This work was supported by Lupus Research Alliance (416805) and NIH R21AI144838 (to MLL), and the Veterans Affairs Merit Review Award (to VPW), ERC StG No 804710 and the Hector Stiftung II gGmbH (to MM).

