## Supplementary material for "Nuclear lamin B is crucial to the nuclear envelope integrity and extracellular trap release in neutrophils": supplment

**Figure S1**. Related to Figure 1

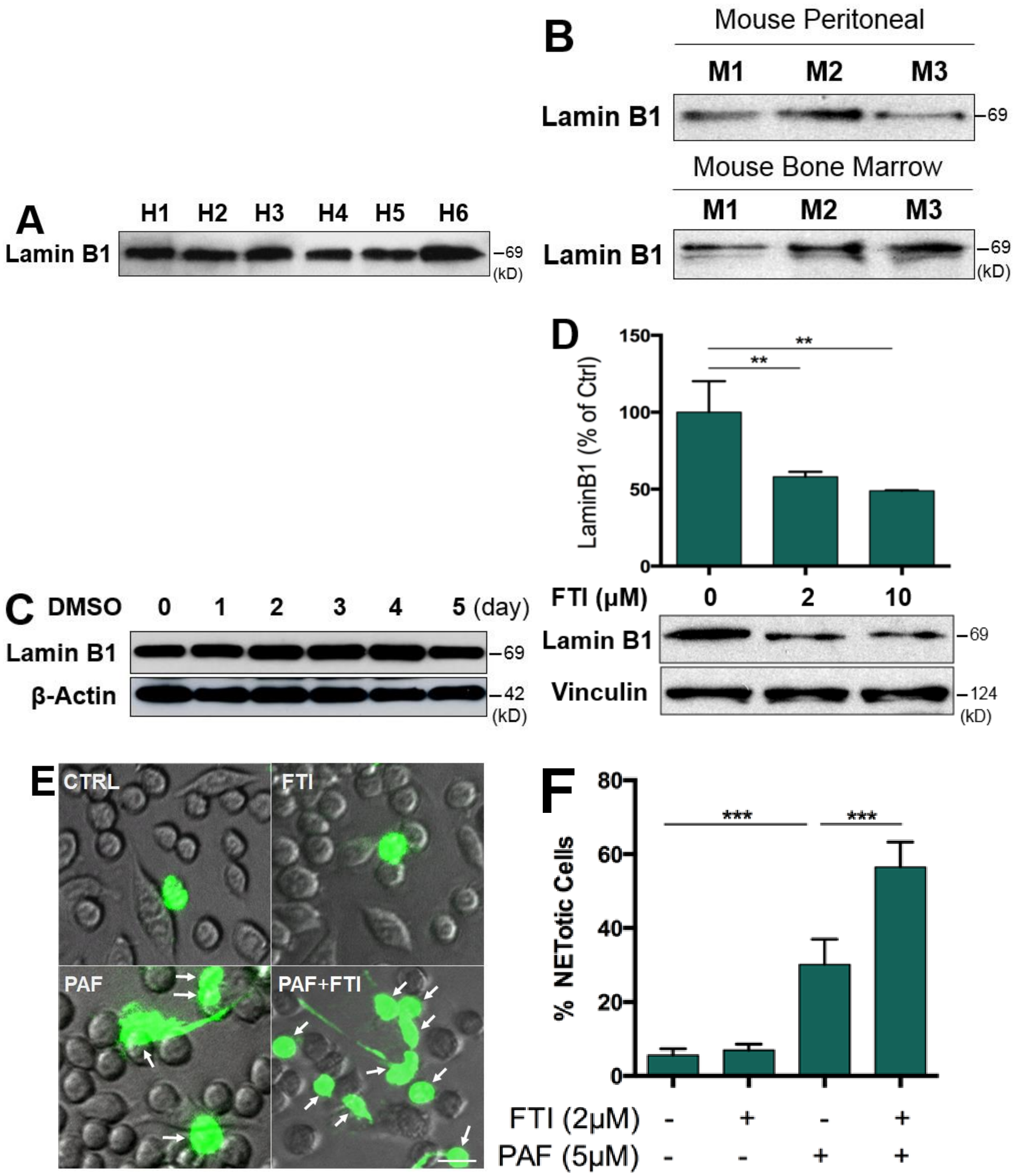


**Figure S1**. **Nuclear lamin B is a substantial component of nuclear envelope that is involved in extracellular traps formation.** (**A**) Representative immunoblot of lamin B expression in primary human polymorphonuclear neutrophils (pPMNs) from six healthy people (H1-H6). (**B**) Immunoblot analyses of lamin B expression in primary mouse polymorphonuclear neutrophils (mPMNs) either elicited from peritoneal cavity or isolated from bone marrow of three representative C57BL/6 WT mice (M1, M2, M3). (**C**) Immunoblot analysis of lamin B expression in human HL-60 cells that were differentiated by 1.2% DMSO from promyelocytic leukemia cells at day 0 to differentiated polymorphonuclear neutrophils (dPMNs) at day 5. β-actin served as loading control. (**D**) Summary and representative immunoblot analysis for the mature lamin B1 expression in RAW264.7 cells that were treated without (0) or with 2 or 10 μM farnesyltransferase inhibitor (FTI) L-744,832 for 48h. (**E,F**) Representative images (E) and summary analysis (F) of extracellular trap formation in RAW264.7 cells that were pretreated without or with 2 μM for 48h, following by treatment or not with 5 μM PAF for 3h, then stained with cell impermeable Sytox Green, without fixation. Fluorescent and phase contrast images were taken by Olympus confocal microscopy, followed by automated quantification of NETs on 5-6 non-overlapping area per well using ImageJ for calculation of % NETotic cells. The white arrows indicate NETotic neutrophils. Scale bars, 20 μm (E). The summary analyses were calculated based on the arbitrary density (B,D) or % NETotic cells by image analysis (E) as compared to their untreated controls, P**<0.01, P***<0.01 between different groups as indicated. Comparisons amongst three or more groups were performed using ANOVA, followed by Student-Newman-Keuls test.

**Figure S2**. Related to Figure 2

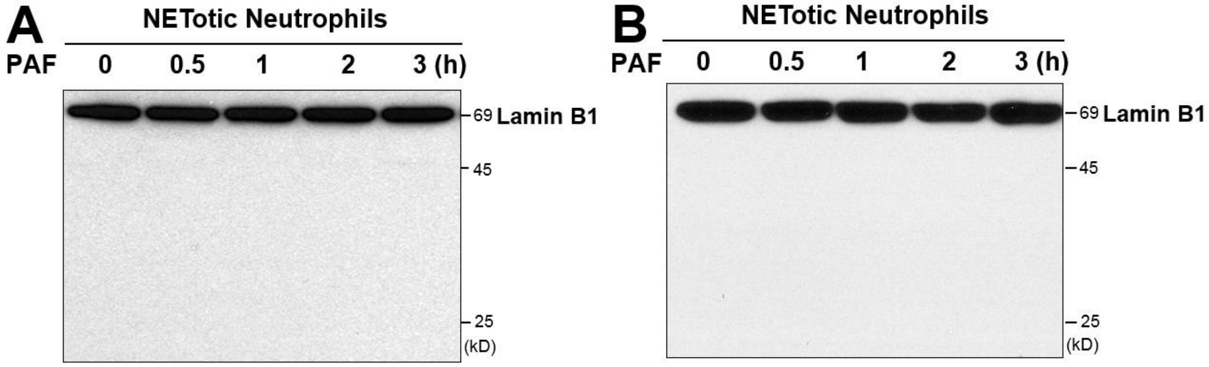


**Figure S2**. **Nuclear lamin B remained as an intact molecule, but not fragmented during PAF-induced NETotic NE rupture.** (**A**) Representative immunoblot image of full-length lanes of lamin B1 in NETotic human dPMNs that were treated with PAF for 0, 0.5, 1, 2, 3h. (**B**) Representative immunoblots of lamin B1 in primary human pPMNs that were treated with PAF (B) for 0, 0.5, 1, 2, 3h during neutrophil NETosis. The anti-human lamin B1 was used in all immunoblots (A,B).

**Figure S3**. Related to Figure 3

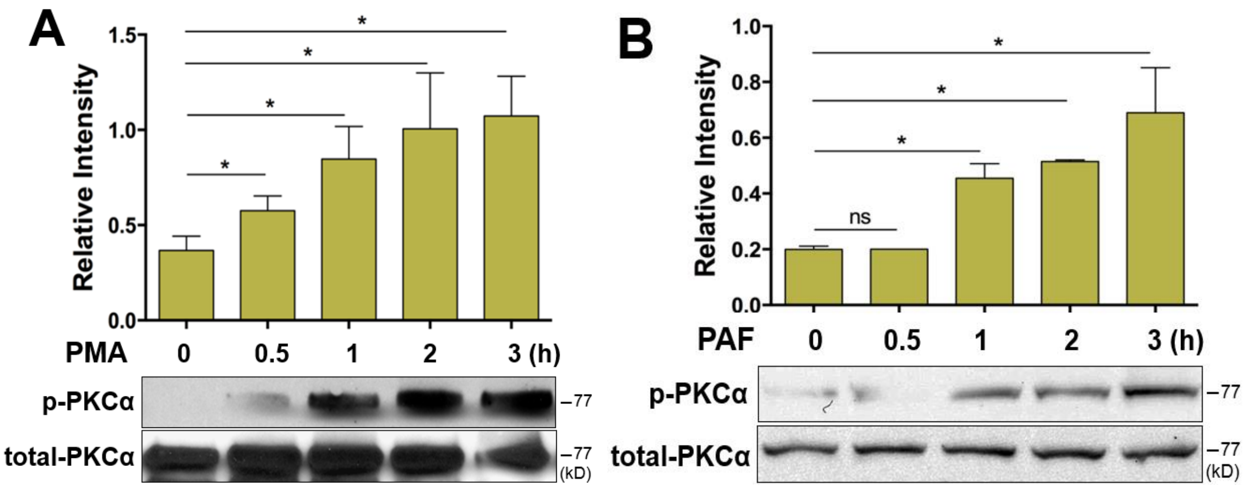


**Figure S3**. **PKCα was phosphorylated in a time-dependent manner during human dPMNs NETosis.** (**A,B)** Representative immunoblots and the summary analyses of total and phosphorylated PKCα (p-PKCα), in human dPMNs that were treated either by PMA (A) or by PAF (B) for 0, 0.5, 1, 2, 3h. Panels A,B were representative immunoblots and their summary analyses that were calculated based on the arbitrary unit from more than 3 independent experiments as compared to their untreated controls, P*<0.05 vs controls. Comparisons amongst three or more groups were analyzed using ANOVA, followed by Student-Newman-Keuls test.

**Figure S4.** Related to Figure 4

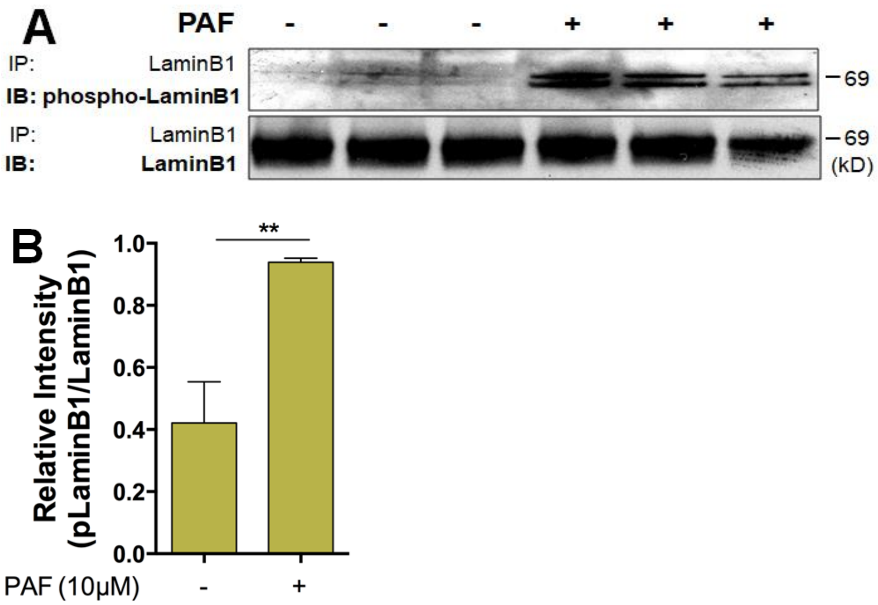


**Figure S4. Nuclear Lamin B was phosphorylated during PAF induced human dPMN NETosis.** (**A,B**) Representative and summary immunoblot (IB) detection of phosphor-lamin B and total lamin B with the lamin B protein purified by immunoprecipitation (IP) with anti-lamin B from human dPMNs that were treated either by PAF (**A,B**) for 0 or 3h. Comparison between two groups was analyzed by student t test.

**Figure S5.** Related to Figure 5

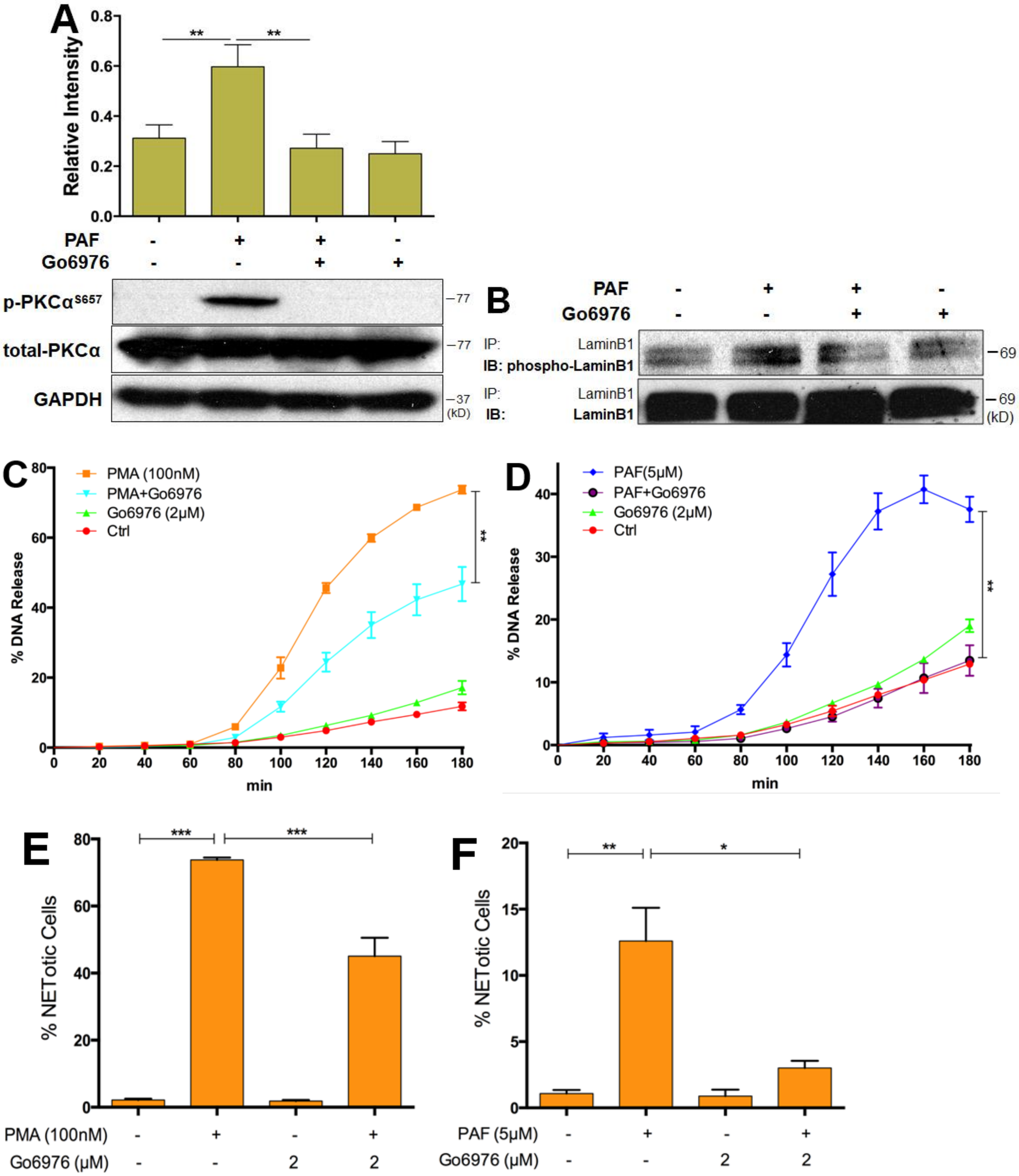

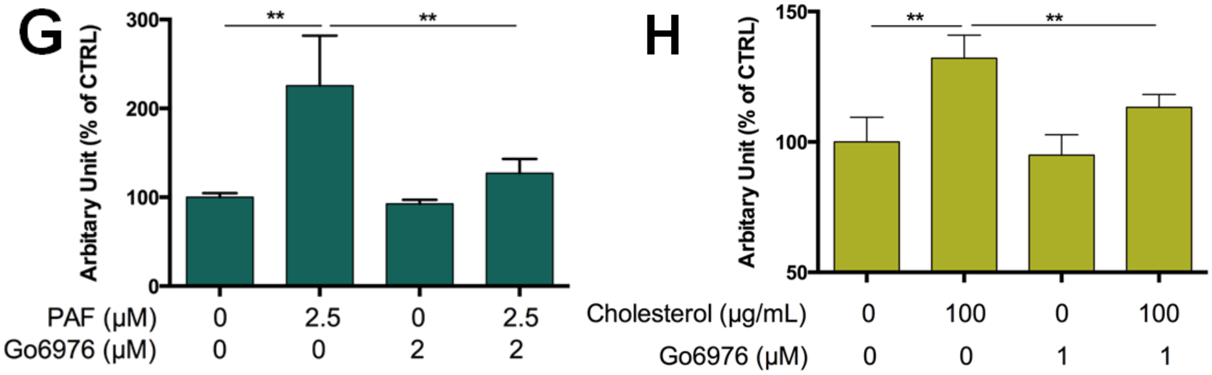


**Figure S5. Inhibition of PKCα phosphorylation significantly reduced extracellular traps formation.** (**A**) Summary and representative immunoblots of the total PKCα and p-PKCα in dPMNs that were pretreated without or with PKC inhibitor Go6976 for 1h, and then treated without or with PAF (B) for 3h. (**B**) Representative immunoblot (IB) detection of the phospho-lamin B and total lamin B with lamin B protein purified by immunoprecipitation (IP) with anti-lamin B from human dPMNs that were pretreated without or with PKC inhibitor Go6976 for 1h, and then treated either by PAF (B) for 3h. (**C,D**) The kinetic analysis of NET-DNA release index were determined by coincubation of primary human pPMNs that were pretreated without or with PKCα inhibitor Go6976 for 1h, and then treated without (control) or with 100 nM PMA (C) or 5 μM PAF (D) in medium containing 1 μM Sytox Green dye for 3h with recording by a microplate reader for every 20 min. The NET-DNA release index was reported in comparison to an assigned value of 100% for the total DNA released by neutrophils lysed by 0.5% (v/v) Triton-X-100. (**E,F**) Summary analysis of PMA (E)- or PAF (F)- induced NETosis in pPMNs that were stimulated without or with 100 nM PMA or 5 μM PAF for 3h, and stained with both cell permeable Syto Red and cell impermeable Sytox Green, without fixation. Images were taken by Olympus confocal microscopy, followed by automated quantification of NETs on 5-6 non-overlapping area per well using ImageJ for calculation of % NETotic cells. (**G**) Summary analyses of extracellular trap formation in RAW264.7 cells that were pretreated without or with PKCα inhibitor Go6976 for 1h, and then treated without or with PAF (G) for 3h. (**H**) Summary analyses of NETosis in primary human dPMNs that were pretreated without or with PKCα inhibitor Go6976 for 1h, and then treated without or with cholesterol loading (H) for 6h. Then the plates (G,H) were analyzed with microplate reader for fluorometric NET quantification. The summary analyses of panels A,C-H were calculated based on the arbitrary unit (A), or NET-DNA release index (C,D), or % NETotic cell (E,F), or arbitrary fluorescent unit (G,H) from more than 3 independent experiments. P*<0.05, P**<0.01, P***<0.001 between groups as indicated. Comparisons amongst three or more groups were analyzed using ANOVA, followed by Student-Newman-Keuls test.

**Figure S6.** Related to Figure 6

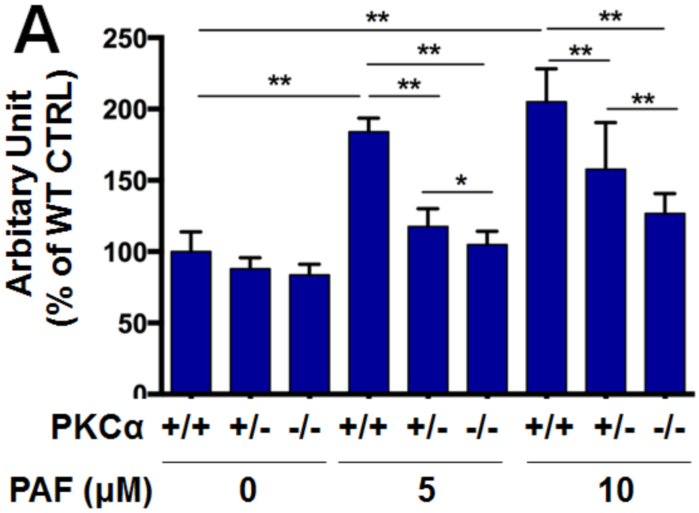


**Figure S6. Genetic deficiency of PKCα attenuates NETosis in bone marrow-derived neutrophils from PKCα deficiency mice.** (**A**) Summary analyses of NETosis in mouse bone marrow mPMNs, from WT, heterozygous, or homozygous PKCα deficient mice, which were treated without, or with either 5 or 10 μM of PAF for 3h, following by fluorometric NET quantification analysis. Panel A is a summary analysis that was calculated based on the arbitrary fluorescent unit from more than 3 independent experiments as compared to their untreated controls, P*<0.05, P**<0.01 between groups as indicated. Comparisons amongst three or more groups were performed using ANOVA, followed by Student-Newman-Keuls test.

**Figure S7.** Related to Figure 7

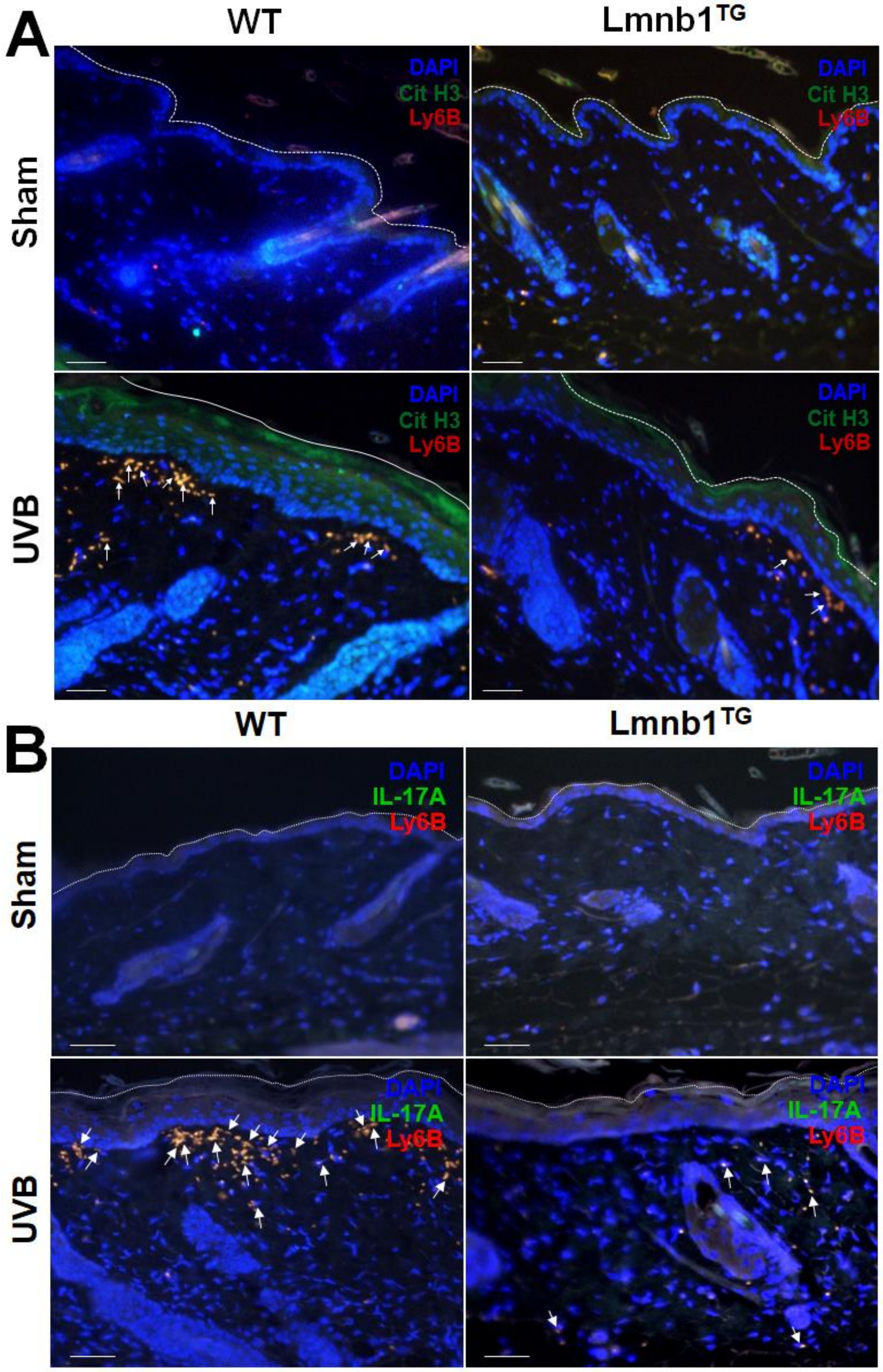

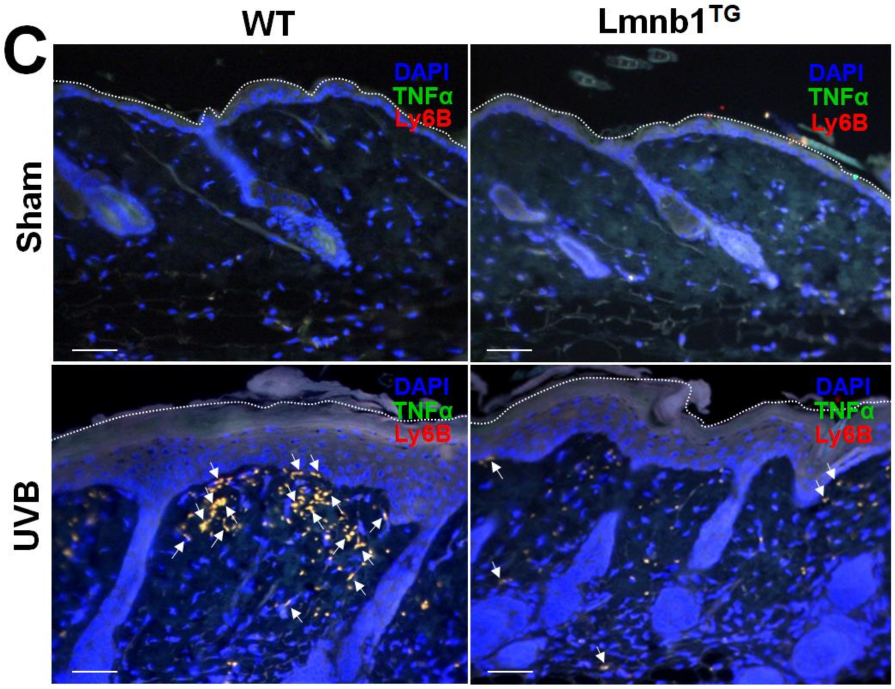


**Figure S7. Lamin B overexpression decreases neutrophil NET release and alleviates NET-associated proinflammatory cytokines accumulation in UVB irradiated skin of lamin B transgenic mice.** (**A,B,C**) Fluorescent staining of the skin tissues from Lmnb1$^{TG}$ mice and their WT littermates that were irradiated or not (sham) by UVB. DNA was stained by DAPI, citrullinated histone H3 was probed by rabbit anti-mouse citrullinated Histone H3, following stained by Alexa Fluor-488 labeled donkey anti-rabbit secondary antibody, while neutrophil surface marker Ly6B was probed by Rat anti-mouse Ly6B Ab following stained by Alexa Fluor-647 conjugated goat anti-rat secondary antibody, IL-17A was probed by Rabbit anti-IL-17A and detected by Alexa Fluor ® 488 conjugated donkey anti-rabbit IgG secondary antibody, FITC labeled Rat anti-mouse TNF-α antibody was used to detect TNF-α. White arrows indicate NETotic neutrophils (A) or NETs with IL-17A (B) or TNF-α (C) display in skin. Scale bar, 100 µm.
